# Counter-balancing X-linked *Mecp2* hypofunction by hyperfunction ameliorates disease features in a model of Rett syndrome: implications for genetic therapies

**DOI:** 10.1101/2024.01.18.576265

**Authors:** Christopher M McGraw, Sirena Soriano, Hao Shuang, Daniel R Connolly, Ali Chahrour, Zhenyu Wu, Agnes J Liang, Yaling Sun, Jianrong Tang, Rodney C Samaco

**Affiliations:** Department of Neurology, Massachusetts General Hospital, Boston, MA, USA; Department of Neurology, Boston Children’s Hospital, Boston, MA, USA; Department of Neurology, Harvard Medical School, Boston, MA, USA; Department of Molecular and Human Genetics, Baylor College of Medicine, Houston, TX, USA; Department of Pediatrics, Section of Neurology, Baylor College of Medicine, Houston, TX, USA; Jan and Dan Duncan Neurological Research Institute, Texas Children’s Hospital, Houston, TX, USA

## Abstract

Treating monogenic neurodevelopmental disorders remains challenging and mostly symptomatic. X-linked disorders affecting women such as the postnatal neurodevelopmental disorder Rett syndrome (caused by mutations in the gene *MECP2*) have additional challenges due to dosage sensitivity and to cellular mosaicism caused by random X-chromosome inactivation. An approach to augment *MECP2* expression from wild-type cells in RTT may be feasible and simpler than gene replacement but has never been tested due to known toxicity of *MECP2* over-expression, as evidenced by the distinct neurological condition known as *MECP2* Duplication Syndrome. Here, using genetic techniques, we find that “counter-balancing” *Mecp2*-null cells in female *Mecp2*-null/+ mice by a complementary population of cells harboring an X-linked transgene associated with 3X normal levels of *MECP2* leads to normalization of multiple whole animal phenotypic outcomes without noticeable toxicity. In addition, *in vivo* LFP recordings demonstrate that counter-balancing *Mecp2* loss-of-function improves select within-region and between-region abnormalities. By comparing the counter-balance approach with an approach based on cell autonomous restoration of MeCP2 using an autosomal transgene expressing 2X normal levels of *MECP2* in all cells (mimicking gene replacement), we identify neurobehavioral and electrographic features best suited for preclinical biomarkers of a therapeutic response to cell autonomous versus non-cell autonomous correction. Notably, these proof-of-concept findings demonstrate how non-cell autonomous suppression of MeCP2 deficiency by boosting overall wild-type MeCP2 levels may be a viable disease-modifying therapy for RTT, with potential implications for genetic-based therapies of monogenic X-linked disorders.

**One Sentence Summary:** In a mouse model of Rett syndrome, counterbalancing mosaic LOF with complementary mosaic GOF improves phenotypic outcome.

## INTRODUCTION

The treatment of many neurodevelopmental conditions remains largely symptomatic at present^1^. Although gene replacement therapy to the central nervous system (CNS) is promising, X-linked conditions in females are further complicated by cellular mosaicism due to random X-chromosome inactivation (XCI). Mosaicism poses a hazard for gene-based interventions if attempts to correct genetic loss-of-function (LoF) in one cell population cause unintended toxic gain-of-function (GoF) in another – an outcome which has been predicted for disorders involving dosage sensitive genes such as *MECP2*.

*MECP2*-related disorders are neurological syndromes associated with perturbations in the X-linked gene *MECP2*^2^, with monoallelic loss-of-function variants causing Rett syndrome (RTT; MIM 312750) and chromosomal duplications spanning *MECP2* causing *MECP2* Duplication Syndrome (MDS; MIM 300260) due to gain-of-function^3,4^. RTT is a neurodevelopmental disability (NDD) affecting girls and women, and is characterized by multiple neuropsychiatric abnormalities including motor incoordination, intellectual disability, epilepsy, and breathing abnormalities^5^. Due to XCI, individuals with RTT express wild-type and mutant *MECP2* in non-overlapping mosaic cell populations^6^. Although gene replacement therapy in RTT mice has shown promising results^7,8^, concerns remain regarding potential risk of gene therapy or other genetic strategies to increase MeCP2 expression in people with RTT^2^.

Due to the exquisite dosage sensitivity of MECP2-related disorders and opposing effects of MeCP2 LoF versus GoF^2,9,10^, we hypothesized that phenotypic abnormalities associated with MeCP2 LoF may be responsive to genetic “counter-balancing” using MeCP2 GoF. If a counter-balancing approach proved fruitful, it would underscore the precise behavioral and electrophysiological abnormalities that could be rescued by non-cell autonomous effects of MeCP2 GoF and establish to what extent genetic correction of MeCP2-null cells is necessary to improve symptoms in RTT. Moreover, by demonstrating the benefit of boosting levels of wild-type MeCP2 in RTT, this proof-of-concept study would have broader implications for genetic-based therapies for X-linked disorders.

Here, we asked whether a strategy to counter-balance cells lacking MeCP2 by over-expressing MeCP2 in the complementary cell population might ameliorate disease, despite both cell populations expressing markedly abnormal levels of MeCP2. We find that multiple neurobehavioral and select electrophysiological abnormalities in adult female *Mecp2*-null mice are rescued purely by non-cell autonomous effects of MECP2 over-expression in the counter-balancing configuration.

## RESULTS

### Counter-balancing a *Mecp2*^null^ allele using an X-linked *MECP2* transgene

To counter-balance MeCP2, we devised a strategy to combine *in trans* the *Mecp2 ^−/+^* null allele^11^ with an *X-linked* human *MECP2* transgene expressing approximately three times the wild-type level of MeCP2 (*hMECP2-*TG3)^9^ to generate female *Mecp2* ^-/TG3^ mice (“NT3”) – these mice possess distinct cell populations either lacking or over-expressing MeCP2 (**Fig. 1A, C**). As a control, we modeled cell autonomous “normalization” of the *Mecp2 ^−/+^* null allele (similar to gene replacement) by generating animals harboring the *Mecp2^−/+^* null allele in combination with an *autosomal* human *MECP2* transgene associated with approximately two-fold over-expression (*hMECP2-*TG1)^9^ to generate female *Mecp2*^−/+^; TG1 mice (“NT1”) – these mice possess cells with roughly normalized or slightly over-expressed levels of MeCP2 (**Fig. 1A, C**). We quantified the cumulative distribution of MeCP2 immunoreactivity (IR) (**Fig. 1D**) from mouse brain cortex by immunofluorescence (**Fig. 1B**). In comparison with female wild-type (WT) mice (**Fig. 1B.i**), we confirm that female *Mecp2* ^−/+^ (“N”) mice possess MeCP2-null neurons which lack MeCP2-IR (**Fig. 1B.ii, *open arrow***) and MeCP2-positive cells (**Fig. 1B.ii, *closed arrow***), each corresponding to ∼50% of quantified cells (**Fig. 1D**). NT1 mice with the autosomal transgene possess only MeCP2-positive neurons (**Fig. 1B.iii**, *open and closed arrows*, respectively) whose intensities are distributed 50:50 within the observed WT range and above that of the highest expressing WT cells (**Fig. 1D**). However, as anticipated, counter-balanced NT3 mice possess MeCP2-null cells (**Fig. 1B.iv, open arrow**) and MeCP2-positive cells (**Fig. 1B.iv, closed arrow**) each corresponding to ∼50% of quantified cells, with the latter population showing MeCP2-IR that is considerably greater on average than any cells from WT or NT1 (**Fig. 1D**). These results confirm that NT3 mice are indeed “counter-balanced”, without ectopic transgene expression or significant skewing of X-inactivation^12^, at least in cortical neurons. Western blot confirmed that both NT3 and NT1 have significantly greater total MeCP2 levels than WT or N mice (**Fig. 1E**). To examine the effect of counter-balancing on transcriptional abnormalities, we evaluated a small set of genes (*Fxyd7, Oprk1, Nxph4,* and *Crh*) known to be sensitive to MeCP2 dosage and altered in the hypothalamus of *Mecp2* mice^10^. NT3 mice show a rescue of deficits in gene expression present in N mice (**Fig. 1F)**. However, NT1 mice are largely similar to NT3, suggesting that either genetic manipulation may normalize aggregate expression of certain MeCP2 sensitive genes in this region.

**Figure 1.**
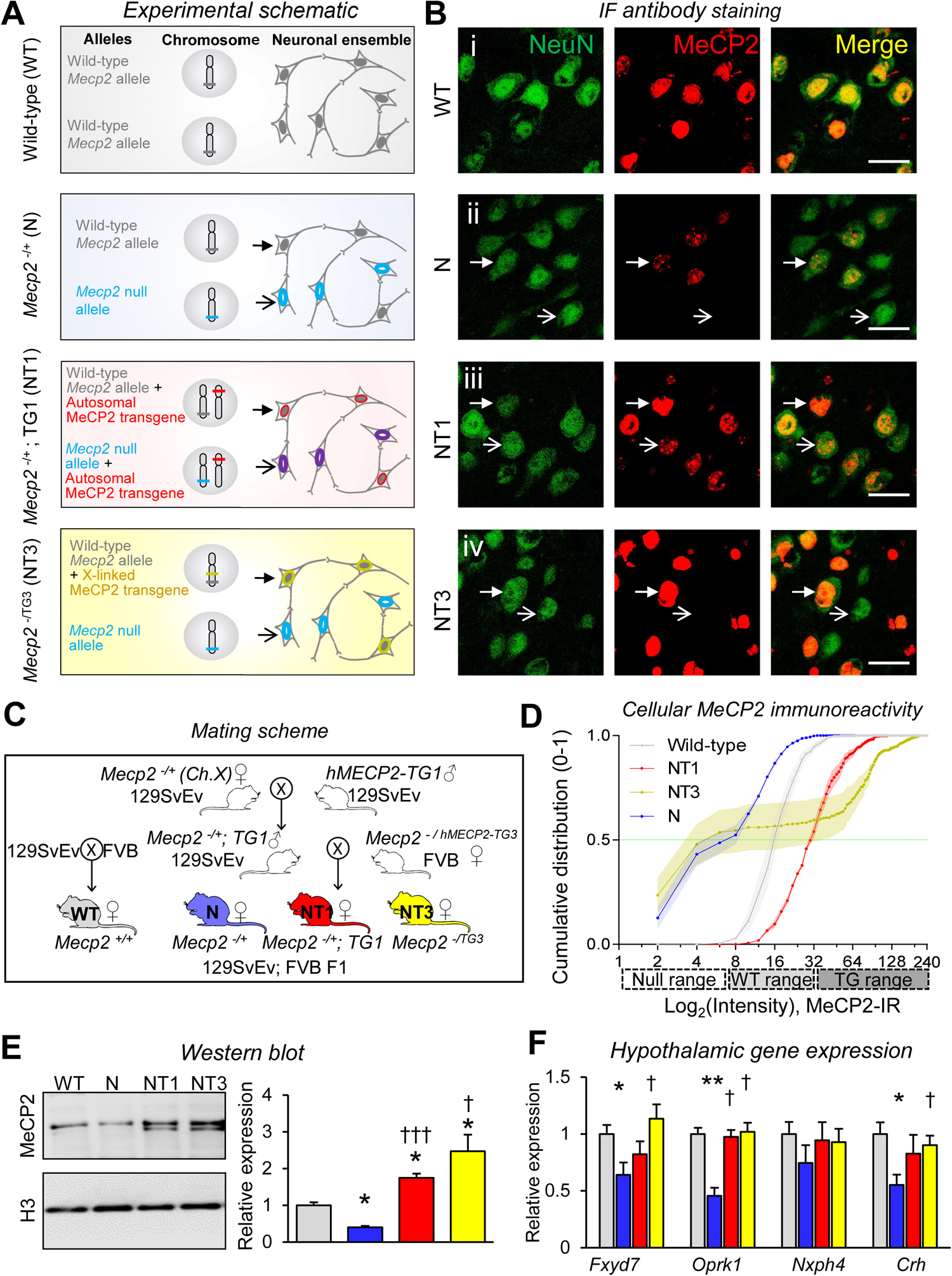
The mouse *Mecp2*^null^ allele can be counter-balanced using the X-linked *MECP2* (TG3) transgene. (**A-B**) Schematic of experimental models used in the study (A) and representative images (B) from each group. For each genotype, panels indicate the Mecp2-related alleles employed, and an illustration of the anticipated mosaic expression pattern, indicated by closed (upper) or open (lower) arrows, adjacent to corresponding confocal immunofluorescence images. Female *Mecp2*^−/+^ (N) heterozygous mice (*blue panel*) possess the *Mecp2*-null allele on one X chromosome and the wild-type *Mecp2* allele on the other. *Mecp2*^−/+^; TG1 (NT1) mice (*red panel*) possess the *Mecp2*-null allele plus an autosomal *MECP2* transgene expressed in all cells. *Mecp2*^-/TG3^ (NT3) mice (*yellow panel*) possess the *Mecp2*-null allele on one X chromosome, counter-balanced by an X-linked *MECP2* transgene on the other. (**B**) Representative confocal immunofluorescence images of cortical neurons corresponding to the schematic diagrams shown in **(A)**. Images show the expected patterns of MeCP2 expression (red, middle), co-localized with neuronal marker, NeuN (green, left). Closed and open arrows indicate cells corresponding to the respective schematized cell from **(A)**, scale bars = 25 µm. (**C)** Schematic of mouse breeding to generate experimental animals. WT are generated from a separate cross of *Mecp2*^+/-^ (129SvEv) x WT (FVB) to maintain hybrid background. **(D)** Cumulative distribution of average MeCP2 immunoreactivity per cortical neuron (quantified from (**B**)) demonstrates close correspondence with the anticipated distributions. Notably, N mice (*blue*) have 50% of cells within the observed “null” range and 50% within “wildtype” range, whereas counter-balanced NT3 (*yellow*) mice have 50% in “null” range, and the remaining 50% are in the “TG” range. Data is average (solid line) +/− sem (ribbon) of proportion of cells at each intensity bin (0-255) along the x-axis. N=3 animals per group. (**E)** Representative western blot (*left*) of MeCP2 immunoreactivity from mouse cortical brain with quantification (*right*) showing relative MeCP2 signal intensity normalized to histone H3. Data represent the mean ± s.e.m., N=4 mice per genotype. *, *p* < 0.05; †††, *p* < 0.01; n.s., not significant). (**F**) Quantitative RT-PCR from hypothalamus demonstrates reduction in gene expression from multiple genes in N mice that are rescued in NT3 mice. Data represent the mean ± sem, N=4-6 mice per genotype. *, *p* < 0.05; †††, *p* < 0.01; n.s., not significant).

### Counter-balancing MeCP2 significantly improves multiple phenotypic and transcriptional abnormalities in female N mice

We tested the effect of counter-balancing MeCP2 on neurobehavioral and physiological outcomes. Interestingly, NT3 mice show rescue or marked improvement for several behavioral phenotypes previously reported in female N mice, including improved anxiety-like behavior (**Fig. 2A**), acoustic startle (**Fig. 2B**), sensorimotor gating (**Fig. 2C**), and contextual fear memory (**Fig. 2D)**. Examination of physiological parameters revealed that breathing abnormalities (apneas)^13–15^ are also rescued in NT3 mice (**Fig. 2E**). Sociability testing in the three chamber apparatus (**Supplementary Fig. 1E**) reveals that N mice have reduced active interest in a partner mouse (vs WT: p<0.01) whereas NT3 may be unaffected (vs WT: n.s., vs N: n.s.). By contrast, open field activity was only rescued in NT1 mice, with female N mice showing increased exploration of an open arena (vs WT: p<0.01; **Supplementary Fig. 1F**) which is not rescued in NT3 mice (vs WT: p<0.01; vs N: n.s.). Similarly, body weight^14^ is elevated in N mice (p<0.05), which persists in NT3 (vs WT: p<0.05; vs N: n.s.), whereas NT1 is reduced even below WT (vs WT: p<0.01, vs N: p<0.01) **(Supplementary Fig. 1B)**. Symptom score^16^ is elevated in N mice, but is similar to NT3 and NT1 mice at this timepoint (**Supplementary Fig. 1C**). Cued fear memory was normal across all animals (**Supplementary Fig. 1A**).

**Figure 2.**
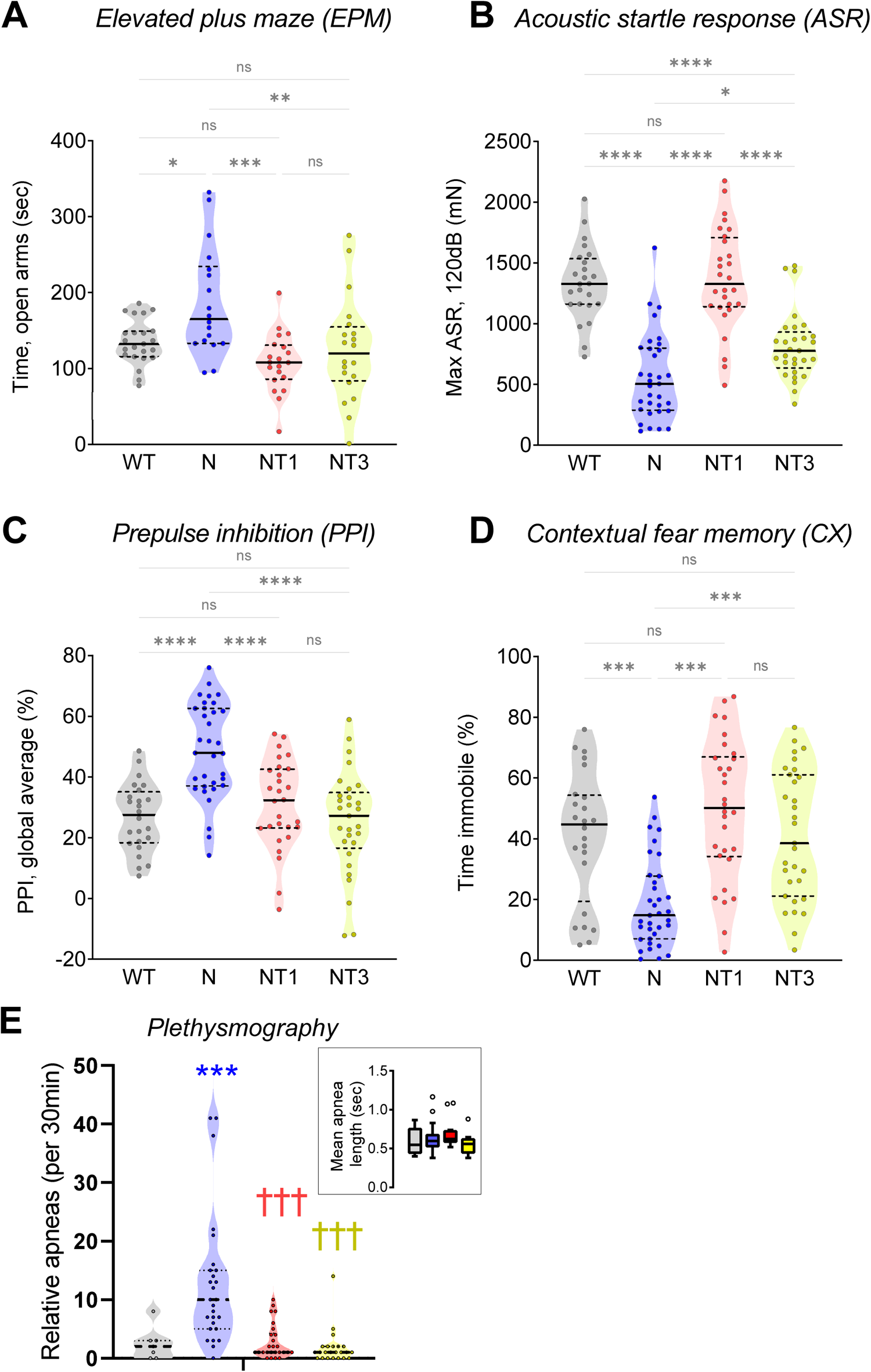
Counter-balancing MeCP2 rescues or improves several behavioral abnormalities observed in N female mice. (**A**) Mecp2^−/+^ (N) mice show abnormally reduced anxiety in the elevated plus maze, which is rescued in NT1 and NT3 mice. (**B**) N mice have reduced acoustic startle, which is rescued in NT1 and significantly improved in NT3 mice. (**C**) N mice have abnormally enhanced prepulse inhibition (PPI) which is rescued in NT1 and NT3 mice. (**D**) N mice have reduced learning/memory in the contextual fear memory assay, which is rescued in NT1 and NT3 mice. Violin plots are mean (solid line) with standard deviation (dotted line), N=24-31 mice per genotype. *, p < 0.05. *p* < 0.01 **(E)** N mice have abnormally high rate of relative apneas^67^, which is rescued in NT1 and NT3 mice. (*Inset*) The mean duration of breaths classified as apneas by the relative apnea index does not differ by genotype. Data are box-and-whisker plots, N= 7 (WT), 28 (N), 26 (NT1) and 23 (NT3) mice per genotype. * indicates difference vs WT; † indicates difference relative to N. For **A-D**, statistics are Tukey’s h.s.d. for a family of 4 comparisons. For **E**, statistics are Wald Chi Square test with Fisher’s LSD test. For each symbol, a single symbol denotes *p* < 0.05; double denotes *p* < 0.01; triple denotes *p* < 0.001, and quadruple denotes *p* <0.0001.

### Counter-balancing MeCP2 is associated with select improvements among multiple within-region electrophysiological abnormalities in female N mice

Given the dependence of normal physiological behavior on proper excitatory/inhibitory (E/I) balance within brain regions^17,18^, we hypothesized that the observed neurobehavioral improvements in NT3 and NT1 mice may be mediated by a common normalization of local “within-region” ^19^ circuit function in behaviorally-relevant brain regions.

To test this hypothesis, we obtained simultaneous *in vivo* electroencephalography (EEG) recordings from four behaviorally-relevant brain regions in the awake animals using depth electrodes implanted in hippocampal CA1 and dentate gyrus (DG) in combination with subdural electrodes over frontal cortex (FC) and somatosensory cortex (SC) (**Fig. 3A**). Assessments of “resting state” EEG activity were performed in unrestrained animals outside of any task paradigm. Mouse movement was tracked by video (see **Methods** and **Supplementary Fig. 2)** and used to ensure that all between-group comparisons were made during similar active versus inactive epochs across animals.

**Figure 3.**
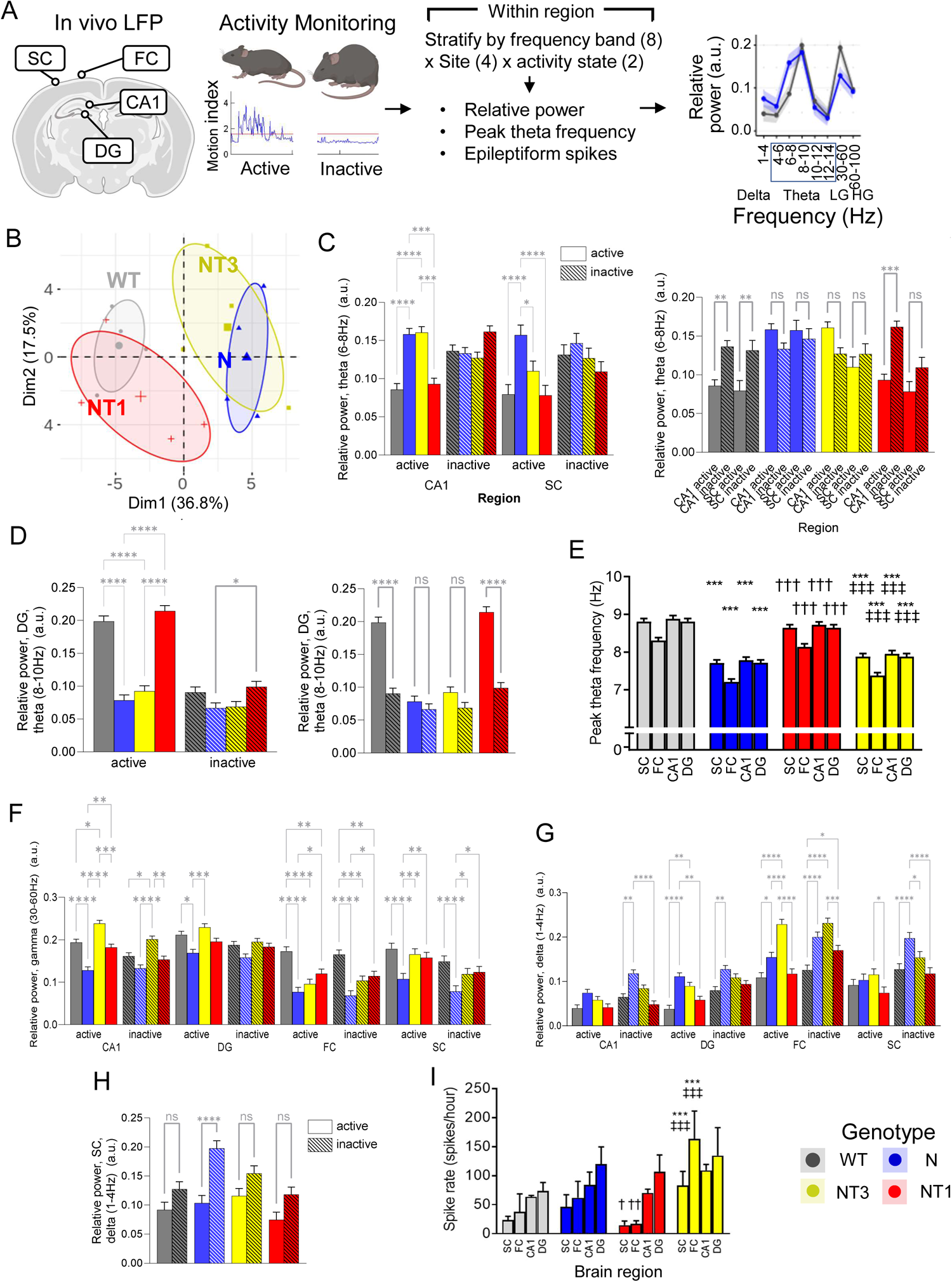
Counter-balancing MeCP2 has select improvements among multiple within-region electrophysiological abnormalities in female N mice. (**A**) Schematic for assessing in vivo LFP from 4 brain regions (CA1, DG, SC, FC) with combined activity monitoring to estimate “within region” measurements of power, peak theta frequency, and epileptiform discharges stratified by activity state. LG, low gamma. HG, high gamma (**B**) PCA of relative band-passed power. Small symbols are individual animals, large symbols are group-wise centroid. Circles are 95% confidence boundaries. (**C**) Abnormal elevations in active-state theta (6-8Hz) power are observed in N and NT3 mice in CA1, but are rescued in NT3 in SC. There are no differences in theta power during inactive state (*left*), and there is loss of the activity-state dependent change in theta power in N and NT3 animals (*right*). (**D**) Abnormal reductions in active state theta (8-10Hz) power in N and NT3 mice in DG (*left*) with loss of activity-state dependent change in theta power in N and NT3 mice (*right*). (**E**) The peak theta frequency is reduced across all regions measured in N and NT3 mice. **(F)** Low gamma power (30-60Hz) is reduced in N mice across regions and activity states, but is improved in NT3 mice in CA1, DG, and SC. **(G)** Abnormally elevated delta power (1-4Hz) in N mice is most prominent in cortical areas during inactive states, which is partly improved in NT3 mice in SC but not FC. (**H**) Activity-state dependent change in delta power in SC is marked in N mice, but not observed in NT3 or other genotypes. (**I**) NT3 mice show elevated rates of high-amplitude spikes measured in cortical areas during active-state. Symbols (*, †, ‡) represent differences relative to WT, to N, and to NT1, respectively. Data are estimated marginal means from linear mixed model ± simulated 95% confidence intervals from fixed effects (N= 4 mice per genotype x 4 sessions x 1-2 hr recording per session) with Tukey adjusted p-values. *, *p* < 0.05; **, *p* < 0.01; ***, *p* < 0.001, ****, *p* <0.0001. CA1, hippocampal region CA1. DG, dentate gyrus. SC, somatosensory cortex. FC, frontal cortex.

We began by assessing relative power. Abnormalities in frequency-specific power are frequently observed in ASD ^20^ including RTT ^21^ and mouse models of RTT ^22–25^ and are theorized to reflect the abnormal neural processing underlying functional impairment in these disorders.

To obtain an unbiased overview, we employed principal component analysis (PCA) --a common data-driven method for reducing high-dimensional spaces by decomposing multivariate data into a smaller set of mutually uncorrelated variables (“principal components”, or PCs, or dimensions), ordered by decreasing percentage of variance explained. We used this method to examine relative band-passed power (8 bands) x 4 brain regions (CA1, DG, SC, FC) x 2 activity levels from individual animals (N=4 per genotype).

Using PCA (**Fig. 3B**), a substantial portion of the total variance in power could be explained with the first two PCs (36.8% and 17.5% respectively). Comparing WT vs N, the 1^st^ PC dimension leads to clear separation between WT and N mice with the 95% confidence boundaries for WT mice and N mice forming non-overlapping clusters. Meanwhile, also within the 1^st^ PC dimension, NT1 mostly overlaps WT, whereas NT3 overlaps only with N mice. Taken together, these findings suggest that counter-balancing MeCP2 does not exert its effect through global normalization of relative EEG power, as assessed by these multivariate techniques. The top factors contributing to separation of WT/NT1 versus N/NT3 in PC1/2 (**Supplementary Fig. 3A**) appear to be related to theta-range power in CA1 and dentate gyrus and gamma range power in cortical areas.

We next used a linear mixed model (LMM) to fit relative band-passed power as a function of brain region, activity, and frequency band with random effects of individual and sessions, extracting the estimated marginal means from the fixed effects for further analysis (see **Methods**).

Focusing on theta-range power, we observe several abnormalities in N mice compared to WT including an abnormally broad theta peak during the active state, due to elevated power in the 6-8Hz range in CA1 (**Fig 3C**, *left*; linear mixed model, adj. p = 8.11E-07) and in SC (**Fig 3C**, *left*; LMM, adj. p = 8.99E-08). These abnormalities persist in CA1 of NT3 mice (versus WT, LMM, adj. p = 3.33E-07; versus N, LMM, adj. p = 0.998) but are rescued in SC (versus WT, LMM, adj. p = 0.115; versus N, LMM, adj. p = 0.00316), and both are rescued in NT1 mice (CA1: versus WT, LMM, adj. p = 0.953; versus N, LMM, adj. p = 1.20E-05; SC: versus WT, LMM, adj. p = 1.0; versus N, LMM, adj. p = 5.44E-08). Nevertheless, the normal activity-state dependent change in theta power is lost in N mice in both CA1 and SC (**Fig 3C**, *right*) and is not rescued in NT3. We also observe abnormal reductions in active-state theta power (8-10Hz) in DG of N mice (**Fig. 3D**, *left*, LMM, adj. p = 9.23E-12) with concomitant loss of the normal activity-dependent increase in theta power (**Fig. 3D**, *right*), which persists in NT3 (versus WT, adj. p = 9.32E-12) but normalizes in NT1 (versus WT, adj. p = 0.656). In addition, peak theta frequency (PTF) is reduced in all brain regions recorded in N mice (**Fig. 3E**), which persists in NT3 mice, but is rescued in NT1 mice. We conclude that abnormalities in theta-range power although numerous in N mice are unchanged in NT3 mice and therefore unlikely to mediate the effect of counter-balancing.

With respect to gamma power, N mice have abnormally *reduced* low gamma power (30-60Hz) in all regions measured during *activity* and in all cortical regions during *inactivity* (**Fig. 3F**). Interestingly, LG power is rescued in several regions of NT3 mice including DG, SC, and CA1, where LG power is elevated slightly higher than WT (active state, versus WT, adj. p< 0.01). However, LG power remains abnormally reduced in FC in NT3, similar to N (**Fig. 3F**). Meanwhile, LG power in NT1 mice follows a pattern of rescue that is similar to NT3, with rescue in DG, SC, and CA1. In FC of NT1 mice, LG power appears incompletely rescued, reduced relative to WT (active state, adj. p < 0.001; inactive state, adj. p < 0.005) but elevated relative to N (active state, adj. p < 0.01; inactive state, adj. p<0.005). Taken together, region-specific improvements in LG power may be a candidate mechanism for phenotypic improvement following counter-balancing.

With respect to delta band power (**Fig. 3G**), N mice have abnormally *elevated* power in DG and FC during *activity* and in all regions during *inactivity*. NT3 mice are mostly similar to N, with elevated power during *activity* in DG and in FC during *activity* and *inactivity* but with some partial improvements during *inactivity* in CA1, DG and SC. NT1 mice are mostly similar to WT except for delta power in FC during *inactivity* which is abnormally elevated. There is a notable activity-state dependent increase in delta power measured from SC in N mice (**Fig. 3H**) which is rescued in NT3 and NT1 mice. In summary, it is difficult to make firm conclusions about the role of delta power in the mechanism of counter-balancing.

We also found that epileptiform discharges – a hallmark of elevated E/I balance^18^ frequently noted in RTT^26^, RTT mice^22,24^, and ASDs ^20^ – were abnormally increased in cortical regions of NT3 mice (**Fig. 3I**; vs WT: p<0.001; vs N:, p<0.001, Fisher LSD) worse than N mice (vs WT, n.s., Fisher LSD), whereas NT1 mice had significantly fewer epileptiform discharges in these regions (vs N, SC, p<0.05; FC, p<0.01), similar to WT. No behavioral seizures were recorded during our observations in any group.

### Counter-balancing MeCP2 is associated with select improvements among multiple abnormalities in inter-regional phase synchronization in female N mice

Inter-regional phase synchronization is postulated to underlie neuronal communication within dynamic networks of the brain ^27–30^, disruption of which has been associated with schizophrenia and ASD ^31^. Given limited improvements observed from “within region” measurements in NT3 mice, we next asked whether normalization of phase synchronization (PS) between brain regions may underlie neurobehavioral improvements of counter-balancing (**Fig. 4A**).

**Figure 4.**
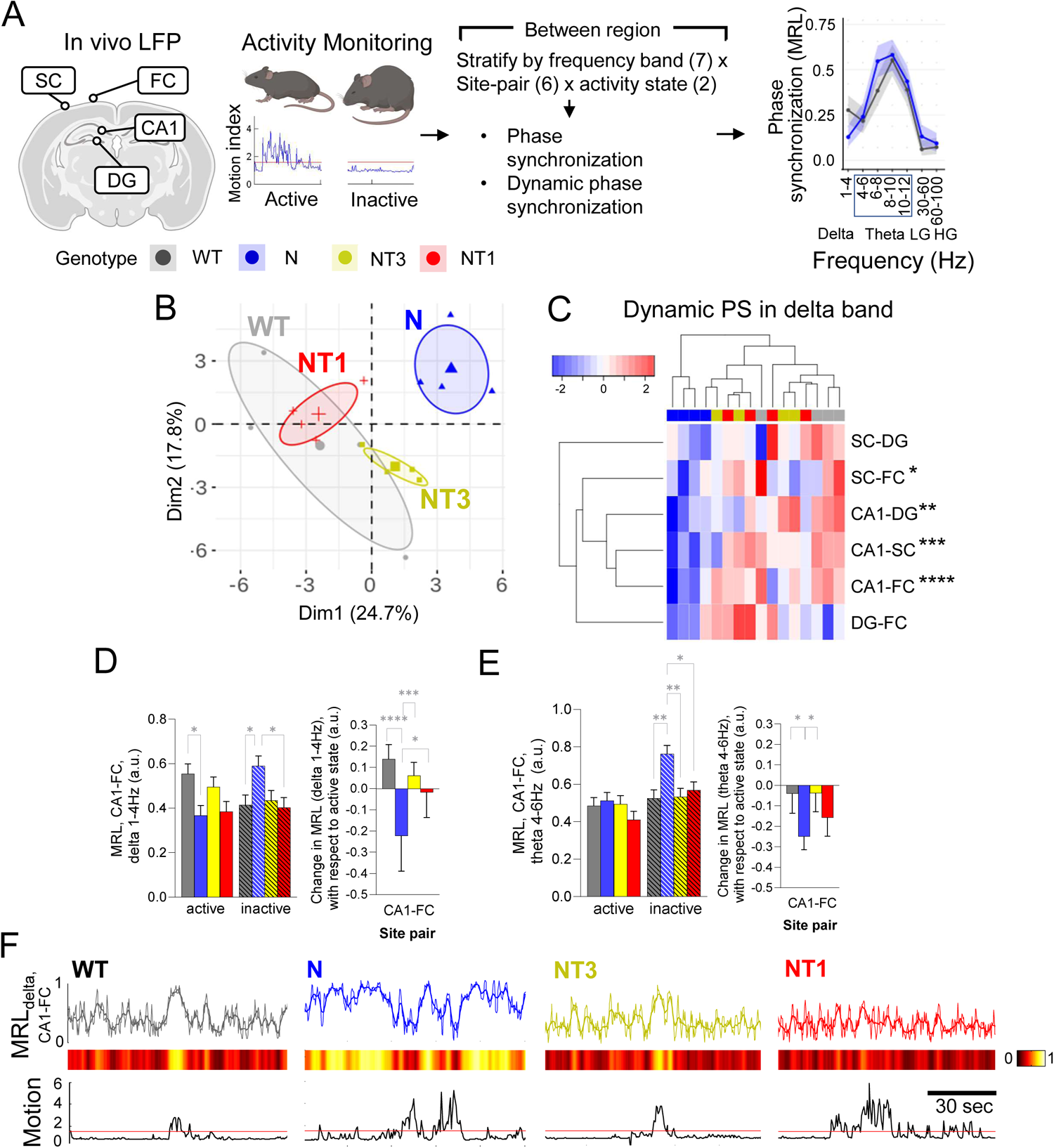
Counter-balancing MeCP2 has select improvements in abnormalities of inter-regional phase synchronization observed in female N mice. (**A**) Schematic for assessing in vivo LFP to estimate “between region” measurements of band-specific phase synchronization within activity states and between activity states (“dynamic phase synchronization”). **(B)** PCA of z-scored dynamic phase synchronization values within frequency bands (7 bands) x 6 brain region pairs (CA1-SC, CA1-FC, CA1-DG, DG-FC, SC-FC, SC-DG) x 2 activity levels for individual animals. Dynamic phase synchronization values are derived from linear mixed models (see Methods) and represent the estimated unit change in phase synchronization between activity levels for each animal. Small symbols are individual animals, large symbols are group-wise centroid. Circles are 95% confidence boundaries. (**C**) Heatmap and hierarchical clustering of dynamic delta-range PS across brain region pairs. Asterisks denote site-pairs for which the groupwise dynamic PS value for N mice is significantly different versus WT. (**D**) Low theta (4-6Hz) PS is abnormally elevated in N mice during the inactive state (*left)* but is rescued in NT3 and NT1. The dynamic PS value is improved in NT3 (*right*). (**E**) Delta (1-4Hz) PS is abnormally reduced during active state and abnormally elevated in inactive state in N mice (*left).* The dynamic PS value is improved in NT3 (*right*). (**F**) Representative time-series from CA1-FC phase synchronization (MRL) in delta 0-4Hz band (*upper*, trace and heatmap) relative to mouse motion index (MI, *lower*). N mice (blue) show striking abnormalities in activity-related dynamics that appear improved in NT3 and NT1 mice. Data are instantaneous estimates of phase synchronization computed on 3-second sliding window (step size, 0.5 seconds) (*thin line*) and smoothed using a 5-second moving average (*thick line*), heatmaps show the moving average of the MRL data. Unless otherwise stated, data are estimated marginal means from linear mixed model ± simulated 95% confidence intervals from fixed effects (N= 4 mice per genotype x 4 sessions x 1-2 hr recording per session) with Tukey adjusted p-values. *, *p* < 0.05; **, *p* < 0.01; ***, *p* < 0.001, ****, *p* <0.0001. CA1, hippocampal region CA1. DG, dentate gyrus. SC, somatosensory cortex. FC, frontal cortex. MRL, mean resultant length.

We assessed PS using the phase-locking value (PLV), or mean resultant length (MRL) of a circular histogram of differences in phase angles as a function of frequency band, site-pair, and activity level. We chose MRL in order to minimize the effect of genotype-related differences in regional power on measurements of PS^32^. To quantify the activity-state dependent change in PS, we extracted the slopes (beta coefficients) corresponding to the fixed effects from a LMM to estimate the unit change in PS between levels of activity within frequency bands and site-pairs for each individual animal (a measure we call ‘dynamic phase synchronization value’, or slopes; see **Methods**).

Using PCA, a substantial portion of the total variance in the dynamic PS values could be explained with the first two PCs (24.7% and 17.8% respectively). In the plane of PC1/2 (**Fig. 4B**), we observed clear separation of N mice from all other genotypes, with NT1 and NT3 both overlapping with WT and not N – consistent with many of our neurobehavioral analyses. Interestingly, NT3 and NT1 continue to cluster apart from one another, suggesting that the two approaches, although similarly distant from N, still differ with respect to dynamic PS in this two-dimensional projection. Reviewing the top contributors to the PC1/2 plane (**Supplementary Fig. 3B**) indicated that variance in dynamic PS in the delta-band contributed to the separation between N vs WT/NT3 while variance in theta-bands contributed to the separation of N/NT3 vs WT/NT1.

Examining the group-wise dynamic delta-range PS (**Fig. 4C**), we observe that N mice show a remarkable inversion of the normal dynamic PS slope across multiple region-pairs (vs WT: CA1_DG, p<0.01; CA1_FC, p<0.0001; CA1_SC, p<0.01; SC_FC, p<0.05). Most notably, between CA1-FC (**Fig. 3D**, *right*), abnormal dynamic PS is normalized in NT3 (vs WT: CA1_FC, n.s.; vs N: CA1_FC, p<0.001) and NT1 mice (vs WT: CA1_FC, n.s.; vs N: CA1_FC, p<0.05). Some additional region-pairs show trends towards normalization in NT3 and NT1 mice (not shown). We also observe that N mice show abnormalities in the low theta 4-6Hz range limited to CA1-FC (**Fig. 4E**) that are corrected in NT3 and NT1, and may suggest a broadening of pathological delta-range coupling into the adjacent theta sub-band during inactivity. Abnormal delta-range dynamic PS in N and its correction in NT3 and NT1 can be appreciated in time-series from representative animals during spontaneous activity (**Fig. 4F**). This abnormality may be a candidate mechanism.

In summary, we demonstrate that abnormalities in inter-regional phase synchronization and their dynamics are widespread in N mice and are substantially improved by MeCP2 counter-balancing (NT3).

## DISCUSSION

This is the first report to our knowledge regarding the pure non-cell autonomous effects of MeCP2 over-expression on the phenotype of female *Mecp2 ^−/+^* heterozygous mice – a configuration we call “counter-balancing”. Contrary to the intuition that counter-balancing would worsen neurological disease, we find that counter-balanced NT3 mice exhibit marked improvements in select behavioral, physiological and electrophysiological domains (**Fig. 5**), which predicts clinical implications for the treatment of RTT.

**Figure 5.**
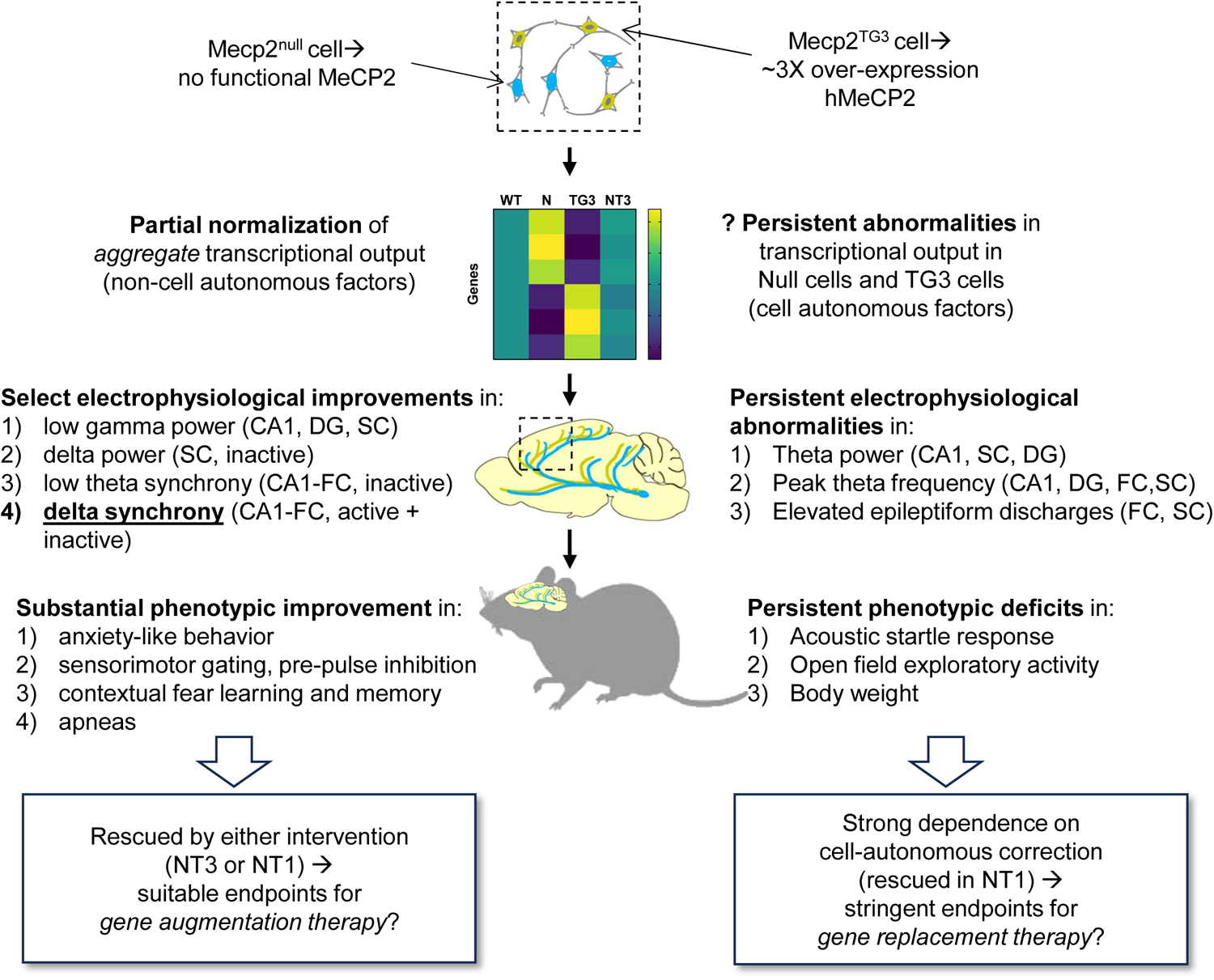
Proposed mechanism of MeCP2 counter-balancing and its impact on preclinical endpoint selection. Abnormalities in female N mice that are rescued in counter-balanced NT3 mice are noted on the *left* while abnormalities that persist in NT3 mice are shown on the *right.* We speculate that 3X over-expression of MeCP2 in 50% of cells (*1^st^ row, left*) leads to partial normalization of transcription (*2^nd^ row, left*), which underlies select electrophysiological improvements (*3^rd^ row, left*), most notably in the dynamics of delta-range synchrony between CA1/FC, leading to substantial phenotypic improvement (*4^th^ row, left*). These endophenotypes may be suitable preclinical endpoints for strategies in N mice that do not correct MeCP2 loss-of-function. Meanwhile, it is speculated that transcriptional abnormalities related to MeCP2 remain abnormal at the single cell level (*1^st^ row, right*), explaining the persistent electrophysiological abnormalities (*3^rd^ row, right*), most notably within-region measures of theta-range power and epileptiform discharges, which underlie the remaining phenotypic deficits (*4th row, right*). These endophenotypes were only corrected in NT1 mice, suggesting a strong dependence on cell-autonomous correction of MeCP2 LoF, and may be stringent preclinical endpoints for gene replacement therapy.

The rescue of hippocampal-dependent contextual fear conditioning (CTX) by counter-balancing is a striking behavioral result. Although phenotypic rescue of CTX has been reported in the context of cell autonomous (CA) correction of *Mecp2* (e.g. with the MECP2 Tg1 allele used in our study^33^), to our knowledge there are no previously reported conditional restoration (CR) experiments demonstrating rescue of CTX. Given rescue of CTX in N mice by a non-cell autonomous (non-CA) mechanism following forniceal DBS^34^, it is possible that the effect of counter-balancing within the basal forebrain might normalize cholinergic transmission, leading to similar rescue of CTX in NT3 mice. Next, the rescue of anxiety on EPM, startle, and PPI fits a pattern reminiscent of the conditional restoration of *Mecp2* in excitatory neurons (*CamkII*-cre^35^). This may suggest that MeCP2 *over-expression* among *half* of glutamatergic neurons might compensate for the remaining MeCP2-null neurons to achieve rescue in NT3 mice, similar to the effect of *Mecp2 normalization* in *all* glutamatergic neurons in *CamkII* CR animals^35^.

Our results also inform the use of female *Mecp2 ^+/−^* mice as preclinical models (**Fig. 5**). As ASR and OFA show stronger dependence on CA correction (i.e. rescued more fully in NT1 mice), this may suggest that these assays could be stringent preclinical endpoints for interventions such as gene replacement. Meanwhile, as EPM, PPI, CTX and apneas are significantly improved by either manipulation, these assays may be suitable endpoints where CA correction is not the goal of treatment or is not able to be achieved.

EEG biomarkers offer a translational bridge between preclinical animal models and human clinical populations, but identifying which of many observed abnormalities are most predictive of phenotypic improvement can be challenging. Importantly, despite differences in mouse background, Mecp2 alleles, and animal numbers versus previous studies, the present study recapitulates multiple within-region EEG abnormalities previously reported in female N mice, including reduced gamma power in frontal cortical areas^36^ and hippocampal CA1^24^, increased delta power in cortical areas^36^, reduced peak-theta frequency^23,24^, and the presence of epileptiform discharges^24,25,36,37^. We also report two previously unreported abnormalities related to theta-band activity in the N mice including abnormal power increase in the 6-8Hz band in CA1 (which may be a manifestation of reduced peak theta frequency), as well as abrogation of activity-state dependent theta power in the DG. We did observe significant elevations in cortical discharges in the counter-balanced mice, though no seizures were observed. We also observed no group-wise differences in the rate of epileptiform discharges assessed in hippocampal CA1 and DG, suggesting that deleterious effects of counterbalancing MeCP2 on hyperexcitability may be brain region-dependent.

The critical lesson of counter-balancing is that very few of these local electrophysiological abnormalities are improved (with the exception of low gamma power) despite significant behavioral improvement, whereas the effects of CA rescue are more extensive. This suggests that EEG biomarkers for preclinical endpoints should be chosen carefully, again based on a desired level of stringency. The extent to which normalization of local gamma power contributes to observed behavioral improvements in the NT3 mice should be addressed in future experiments.

The mechanisms of inter-regional communication within brain networks remains an area of active interest in health and disease, particularly in neurodevelopmental disorders. To the best of our knowledge, there are no published studies evaluating between-region measures such as inter-regional phase synchronization in the Mecp2 model during task-dependent or resting states, and there are similarly limited reports in related mouse models of neurodevelopmental disorders. We report abnormal dynamics of phase synchronization among deep (CA1, DG) and cortical areas (FC, SC) which in aggregate are sufficient to separate N mice from WT and NT3/NT1 mice following dimensionality reduction using PCA. Among these measurements, dynamic phase synchronization in the delta range particularly between CA1-FC appears dramatically abnormal in N mice, and is rescued both in NT1 and NT3 mice. This phenomenon is distinct from a previously reported abnormality in dynamics of cortical delta *power* ^25^. Delta-band phase synchronization has been observed across distant sites in brainstem, FC, and CA1 ^38^ and may have a role in homeostatic processing^39^. The dynamics of delta-range phase synchronization with activity have not been previously reported, but may contribute to the observed behavioral improvements of counter-balancing.

Although we have not definitively demonstrated the mechanism of counter-balancing, we have excluded many plausible possibilities and highlighted several avenues for additional investigation. Our speculative model (**Fig. 5**) is that electrophysiological and phenotypic improvement is due to mechanisms sensitive to *partial normalization of aggregate transcriptional output*. There are few candidate mechanisms for which non-cell autonomous correction on such a scale is neurobiologically plausible, therefore *normalization of non-synaptic extracellular neurotransmitter dynamics* (for example, by normalizing the aggregate output of projection neurotransmitter systems from brainstem and basal forebrain during different activity states) may be the most parsimonious. This possibility is supported by several disparate lines of evidence. First, the known physiology of projection neurotransmitter (NT) systems providing tonic and phasic adjustments in regional levels of non-synaptic NTs that govern behavioral states ^40,41^, regulate expression of hippocampal memory formation^42^, and between-region neural synchrony ^43^, all of which appear improved in the NT3 mice. Normalization of extra-synaptic NT would naturally be non-CA as it does not depend on CA normalization of Mecp2 in any specific brain region. Second, the effects of MeCP2 LOF vs GOF on transcriptional output have been well-documented^10^, with many reports demonstrating opposing changes in common sets of altered genes in *Mecp2*-KO vs WT compared to *Mecp2*-TG1 vs WT. Opposite changes in transcription provide an intuitive explanation for why the core pathophysiology of RTT might be complementary to the pathophysiology of MeCP2 over-expression, as the portion of projection neurons with reduced transcriptional output due to *Mecp2* deficiency could be balanced by the enhanced output of the complementary set of MeCP2 over-expressing neurons in the NT3 mice. Third, there is a long history of altered biogenic amines in RTT and mouse models of MECP2 disorders extending back to the earliest attempts to understand the disorder^44^. Multiple modern reports have corroborated reductions of multiple neurotransmitter systems in RTT, including reduced neuroaminergic metabolites in CSF of RTT patients^45^ correlating with severity of MECP2 mutation type; and in brains of male mice lacking *Mecp2*^45,46^ and from metabolomic studies of *Mecp2* ^-/y^ mouse cortex which disclosed significant reductions in acetylcholine, dopamine, serotonin, and related metabolites^44^. These NTs are derived from projection neurotransmitter nuclei^47^ in the brainstem and basal forebrain which project to widespread areas of the brain. The causal role of altered NT levels or projection NT systems in RTT pathophysiology has been demonstrated in multiple ways, including: CKO of *Mecp2* in TH-positive or PET1-positive cell populations^45^ recapitulating the reductions in neuroamingergic metabolites, suggesting they are not merely a consequence of disease severity; phenotypic improvements following CR in catecholaminergic neurons^23^, forniceal DBS^34,48^ in N mice, and treatment with beta-adrenergic agonists in N mice^49^. There are other possibilities, and further experiments are needed to dissect the neuroanatomic and cell-specific requirements of the counter-balancing effect.

Our findings have several immediate implications for treating RTT. *First*, whether over-expression of MeCP2 is pathological appears to depend on the molecular and neuroanatomical context, with counter-balancing demonstrating how MeCP2 over-expression appears to suppress features of disease related to MeCP2 deficiency when combined non-cell autonomously. Although we did observe increased epileptiform discharges in NT3 mice, this may be region-dependent and further understanding of counter-balancing may enable more precise deployment of the technique to minimize this potential harm. *Second*, preclinical neurobehavioral and electrophysiological endpoints for candidate treatments for RTT should be selected carefully, as many appear not to correlate with otherwise worthwhile symptomatic improvement. Our findings help to stratify which phenotypes in N mice may be most useful. *Third*, viable disease-modifying therapies for RTT may not require cell autonomous correction of MeCP2 levels. It is commonly assumed that the most promising disease-modifying treatment of RTT will be gene replacement^50,51^, and that this therapy must normalize MeCP2 function in the mosaic population of cells harboring loss-of-function alleles while simultaneously minimizing toxic effects of MeCP2 over-expression in unaffected cells. However, both of these assumptions are challenged by the present study. Our findings suggest that strategies to increase total levels of wildtype MeCP2 in individuals with RTT may be tolerable and efficacious for improving critical features of the disorder such as learning/memory. This understanding opens the door to alternative pharmacological and genetic strategies to augment levels of endogenous MeCP2^52^ which may hold more therapeutic promise than previously assumed.

In conclusion, our approach provides a powerful framework for further investigations into the mechanisms by which MeCP2 over-expression suppresses MeCP2 hypofunction non-cell autonomously, which may enable new therapeutic avenues for Rett and possibly other dosage-sensitive X-linked neurodevelopmental disorders.

## METHODS

### Generation of mouse models

Animals for the reported studies were maintained on a 12 h light:12 h dark cycle with standard mouse chow and water ad libitum according to IACUC standards. Three MeCP2 alleles were used for the creation of test progeny: the *Mecp2*^tm1.1Bird^ allele^11^ (abbreviated “Null” or “N” in this study) which lacks exons 3 and 4 producing a null allele of the endogenous X-linked *Mecp2* locus; the *Mecp2*-TG1 allele^9^ which harbors an autosomal insertion of the human *MECP2* transgene (abbreviated “TG1” or “T1” in this study), and the *Mecp2*^+/TG3^ allele^9^ which harbors an X-linked human *MECP2* transgene (abbreviated “TG3” or “T3” in this study); the TG1 and TG3 models express two and three-fold increased endogenous MeCP2 protein; respectively. Null and TG1 mice were maintained on an a pure 129S6/SvEvTac background (129SvEV), and TG3 mice were maintained on a pure FVBN/J background (FVB). To generate test progeny, female N mice and male TG1 mice were mated to generate male *Mecp2*^-/y^; TG1 (NT1) mice; these male NT1 mice were mated with female TG3 mice to generate female N, NT1, and NT3 littermates. Based on prior published studies showing effects of genetic strain background in *Mecp2* mouse models^14^, the two generation breeding scheme ensured all test mice were of an F1 hybrid genetic strain background. Age- and sex-matched wild-type controls with the same 129SvEv;FVB F1 hybrid background were generated by a separate cross. Animals were genotyping using a quantitative PCR method (see **Supplementary Methods and Supplementary Fig. 4**).

### Behavioral analysis

Mice were tested at 16-20 weeks of age, unless otherwise indicated. Only female mice were used for analysis, due to the nature of the experiment involving the X-linked gene *Mecp2* and the X-linked MeCP2 transgene. Standard protocols were used to characterize behavioral abnormalities in mice, essentially as reported in our previous work^14^, and described in more detail below. Statistical analysis was performed using SPSS (version 19). Data from elevated plus maze, open field, acoustic startle, PPI, and fear conditioning were analyzed using a one-way analysis of variance (ANOVA) with genotype as a factor followed by Fisher’s LSD test. Weight was analyzed using a one-way ANOVA with repeated measures.

#### Elevated plus maze

Animals were habituated to the test room (150 lux, 60 dB white noise) for 30 min. After the habituation period, animals were placed in the center of a maze consisting of two arms (each arm 25 × 7.5 cm) enclosed by approximately 15 cm high walls, and two open arms (each arm 25 × 7.5 cm, with a raised 0.5 cm lip around the edges) elevated 50 cm above ground level; the arms of the maze were equidistant from the center platform. The amount of time animals spent in the open arms, the number of arm entries and the total distance traveled were recorded for 10 min using a camera and detection software (ANY-maze video tracking system, Stoelting Co., Wood Dale, IL, USA).

#### Acoustic startle response (ASR) and pre-pulse inhibition (PPI) of the ASR

The ASR and PPI of the ASR were measured as previously described^14^. To determine ASR to sound ranging from 78 to 118 dB, a total of 13 sounds (0 and 78–118 dB in 4 dB increments) were randomly presented, and activity was recorded as previously described using a startle chamber for mice (SR-Lab, San Diego Instruments, San Diego, CA, USA). These sound stimuli were also presented in random order with or without a prepulse stimulus 4, 8, or 12 dB above background. The response of the mouse to the sound stimuli was recorded by accelerometer to assess startle response, and inhibition of the startle response by the presence of a prepulse stimulus.

#### Conditioned Fear – Contextual (CX) and Cued Fear (CS) Memory

CX and CS fear conditioning was performed as previously described^14^ with a shock intensity level of 0.7 mA. Animals are first exposed to two successive rounds of a conditioned stimulus-unconditioned stimulus pair (80 dB sound, followed by a mild foot shock), then are tested 24 h later for both contextual memory, which is hippocampal dependent, and cued memory, which is hippocampal and amygdalar dependent. Footshock control tests were performed as previously described^14^ and were similar between groups.

#### Open field activity

Animals were habituated to the test room (150 lux, 60 dB white noise) for 30 min. After the habituation period, animals were placed in the center of a 40 × 40 × 30-cm chamber equipped with photobeams (Accuscan, Columbus, OH, USA) to record activity during a 30 min test period.

#### Three chamber test of sociability

Novel partner mice were placed within a small wire cage as previously described^14^ to habituate them to their test environment, these animals were placed randomly in either the left or right test chamber for ∼30 min to 1 h per day for at least 2 consecutive days prior to the actual test day. On the day of testing, test animals were first habituated to the test room for 30 min (150 lux, 60 dB). A test for side preference was then performed by placing the test animals in the center of the three-chamber apparatus. The entries to either the left or right chambers were unobstructed during the side preference test, allowing animals to freely explore and spend time in all three chambers. The time spent in each chamber was measured for 10 min. Next, during the social approach test (novel mouse versus object), the amount of time test animals spent in all three chambers and interacting with either a novel mouse or object was measured. Novel partner mice were randomly assigned in either the left or right chamber to avoid the potential of a side bias; this was designed in a manner to expose half of each test subject genotype to a novel mouse in the left chamber, and the remaining half of each test subject genotype to a novel mouse in the right chamber.

#### Symptom score

Mice were visually evaluated across 6 symptom domains based on a method modified from Guy et. al. 2007^16,53^. Briefly, domain-specific subscores (0 = not present; 1= mild; 2 = severe) were assigned for mouse mobility, gait, tremor, breathing, general condition, and hindlimb clasping. The sum of subscores (0-12) is presented as the total symptom score for each mouse.

### Plethysmography

Studies of mouse breathing were conducted as described in previous work^14^. Briefly, mice were allowed to acclimate for 20 min in unrestrained whole-body plethysmography chambers (Buxco, Wilmington, NC, USA) with a continuous flow of fresh air, and baseline breathing was then recorded for 30 min. Breath waveforms were captured using Biosystem XA software (Buxco, USA) and processed in Matlab using custom code to segment waveforms by a zero-crossing method and estimate time exhaled (TE). Relative apneas^14^ were classified using a moving threshold for each mouse defined by global and local properties of the breathing activity; relative apneas were defined as TEs exceeding both the average TE for the recording period by 3-fold and the local average TE (within a 5-breath window, centered on the breath) by 2-fold. Apnea count data were not normally distributed and heavily zero-inflated, therefore a generalized linear model (GzLM) with negative binomial distribution and log link was used in SPSS. The model was minimally over-dispersed (deviance/df = 1.852). Statistical testing was performed by Wald Chi Square test with Fisher’s LSD post-hoc test.

### Western blot, immunofluorescence, transcriptional analysis are provided in Supplementary Methods

#### Electrophysiology

Mice at 10 months of age were anaesthetized with isoflurane and mounted in a stereotaxic frame. Under aseptic conditions, each mouse was surgically implanted with four recording electrodes, two of which were aimed at the subdural space of right frontal cortex, right somatosensory cortex (Teflon-coated silver wire, 125 μm in diameter) and two of which were aimed at the ipsilateral hippocampus and dentate gyrus. The reference electrode was then positioned in the occipital region of the skull. All electrode wires were attached to a miniature connector (Harwin Connector). After 3–5 days of post-surgical recovery, EEG activity (filtered between 0.5 and 5 kHz, sampled at 2 kHz) and video were simultaneously recorded in freely moving mice without anesthesia across multiple days. Mice were recorded for 60 min (session 1), 120min (session 2; 4 days after session 1), 120min (session 3; ∼3 months after session 2), and 120min (session 4; 4 days after session 3). Signals from electrodes were acquired and amplified x1000 (A-M systems, 1700 Amplifier) and digitally sampled at 2 kHz (Molecular Devices, 1440A Digitizer and pClamp10) ^54^. Following experimentation, an anodal current (30 mA, 10 s) was passed through the tungsten wires for identification of the electrode placements. Mouse brains were harvested per routine and the position of electrodes implanted in CA1 and DG confirmed by coronal sectioning using standard histological techniques^55^ For all mice examined, CA1 electrodes were either in CA1 or in adjacent corpus callosum in all mice, and DG electrodes were located in DG (**Supplementary Fig. 5**).

#### Video analysis to quantify mouse motion

A quantitative motion index (MI) for each mouse was calculated from video recordings using Matlab. Briefly, implanted mice were tested in batches of four while held in separate transparent chambers while engaged in unrestrained naturalistic behavior. Video of mice (four per frame) was acquired synchronized with EEG recording in MTS format using a Sony Camcorder. Video was converted to MP4 using MKVToMP4 and further processed in Matlab by de-interlacing; down-sampling to 1 frame per second; extraction of subframes containing individual mice; background subtraction using morphological opening; and histogram equalization. For each subframe, the MI was calculated as a continuous time series representing the sum of the absolute difference in pixel intensity between consecutive subframes with respect to time. The resulting MI represents the average pixel-wise absolute intensity change per subframe over time and tracks with mouse movement (**Supplementary Fig. 2**). A subset of video data was manually scored by two reviewers blinded to genotype to indicate when mice were actively moving or not moving as a comparison, which demonstrated the computerized method had greater accuracy (data not shown). A distinct mode in the MI was seen between 0.7-1.0 during mouse inactivity across mice (data not shown), therefore a threshold of MI >1.6 was chosen empirically to represent activity. Video artifacts (such as accidental movement of the camera, change in lighting) were excluded by excluding any portion of the time-series with MI >5 from analysis.

#### Analysis of electrophysiological signals

Analysis and statistical evaluations of electrophysiological recordings were conducted using MATLAB R2013b and R (2013), using custom code ^54,56–62^. Briefly, signals were obtained as continuous time series from subdural and depth electrodes at 2kHz. Signals were zero-meaned. Data was notch-filtered at 60Hz using a Butterworth IIR filter except for analysis of phase synchronization for which phase-preserving finite impulse response (FIR) filters were employed (see section below). Unless otherwise stated, all analyses were performed on concatenated epochs of time series data according to behavioral activity state.

##### Power spectral density (PSD)

Although some authors have recently reported rodent EEG using nomenclature from human EEG (i.e. referring to 4-8Hz as theta, 8-12Hz as alpha, and 12-30Hz as beta^36,63^), we choose to refer to the entire range from 4-12Hz as theta^64,65^, and we do not make reference to alpha or beta ranges. PSD was estimated using Welch’s averaged modified periodogram method (Matlab function *pwelch*) and power within specified frequency bands was derived by integration of the PSD in the following ranges: delta (1-4 Hz), low gamma (30-60 Hz), and high gamma (60-100 Hz). For analysis of theta, we computed power within narrow sub-bands (2-Hz wide) spanning the conventional theta-band, namely: 4-6 Hz, 6-8 Hz, 8-10 Hz, 10-12 Hz, and 12-14 Hz. To avoid confusion, we refer to these sub-bands by their corresponding frequency ranges or letter designations (A-E). *Relative power* was obtained by normalizing band-specific power (sum of power within corresponding frequency bands) by total calculated power from 0-200Hz.

Statistical analysis of relative power (raw data, **Supplementary Fig. 6**) was performed using a linear mixed model (LMM) in R (*lmer* function, package *lme4* and fit using restricted maximum likelihood (REML) with fixed effects of genotype, activity state, frequency band, and electrode site with random intercepts of session and subject. The model formula was: `Power`_relative_ `∼ activity * geno * freqband * sites + (1|session) + (1|idNum).` Appropriate model diagnostics including QQ plots were visually inspected by group. Statistical inference was performed using Tukey contrasts between levels of genotype within FREQ X SITE X ACTIVITY, or between levels of activity within GENO X FREQ X SITE using emmeans, lmerTest, and pairs.emmGrid, with the Kenward-Roger approximation of degrees-of-freedom. P-values from this method are adjusted by the Tukey HSD test for a family of 4 estimates (genotype) with alpha = 0.05. Data is the EMM +/− 95% parametric confidence intervals computed from the fixed effects estimated by the linear mixed model.

##### Principal component analysis

Data were preprocessed by averaging across sessions within individual animals and z-scoring within site or site-pair x frequency band combination. PCA was performed on the preprocessed data using the *prcomp()* function, and visualized using *fviz_pca_ind()* and *fviz_pca_var()* from the *factoextra* library. The contribution score for each variable in two dimensions is calculated based on its average contribution to each dimension, according to: contrib = [(C1 * Eig1) + (C2 * Eig2)]/(Eig1+Eig2), where C1 and C2 are the contributions of the variable to PC dimension 1 and 2, and Eig1 and Eig2 are the eigenvalues of PC dimension 1 and 2.

*Peak theta frequency* was identified for each brain region in active mice by lookup of the frequency (Hz) corresponding to the local maximum of PSD within the theta range (4-12Hz). Statistical analysis was performed using a LMM followed by a simultaneous hypothesis test using pairwise Tukey contrasts between genotypes within levels of site.

##### Epileptiform discharges

Abnormal epileptiform spikes (sharp positive or negative deflections with amplitudes exceeding three times the baseline and lasting 25–100 msec) were detected using an automated zero-crossing method implemented in Matlab and confirmed by visual inspection by two blinded reviewers according to previously published protocols with slight modification^61^. No electrographic or clinical seizures were observed during two hours of analysis. Count data for each animal was pooled from two 120 minute sessions and analysis limited to active periods (MI>1.6). Data were not normally distributed; a generalized linear model (GzLM) with a negative binomial distribution and a log link function (formula: `spikeCount ∼ GENO X SITE + totalTime)` was employed. Multiple comparison testing was performed in R as a simultaneous hypothesis test on the difference in means between levels of GENO within levels of the factor SITE using Tukey contrasts.

*Phase synchronization* between brain regions (*site_x_, site_y_*) was calculated as the mean resultant length (MRL) of the circular distribution of phase differences (*ϕsite_x_* – *ϕsite_y_*) within frequency bands^32^. Briefly, signals were band-pass filtered using two-way least squares FIR filtering as implemented in *eegfilt* ^58^ and phase angles were extracted using the Hilbert transformation. MRL was calculated as the weighted sum of cosine and sine of the phase differences, and is defined from (0,1). Estimates of instantaneous phase synchronization for **Fig. 4F** were obtained by performing the computations as above using a 3-second sliding window translated by 0.5 sec increments (shown as *thin line*) and smoothed using a moving average of 5 seconds (*thick line*). Corresponding heatmaps show the moving average of the MRL data.

For statistical analysis, the arcsine transformation^66^ was applied to MRL data, *p*, to obtain 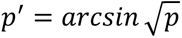 before fitting a LMM with REML (raw data, **Supplementary Fig. 7**) using *lmer* and the following model formula: `asin(sqrt(MRL)) ∼ activity * geno * freqband * compSites + (1|session) + (1|idNum).` Appropriate model diagnostics including QQ plots were visually inspected by group. Statistical inference was performed using Tukey contrasts between levels of genotype within FREQ X COMPSITE X ACTIVITY, or between levels of activity within GENO X FREQ X COMPSITE using emmeans, lmerTest, and pairs.emmGrid, with the Kenward-Roger approximation of degrees-of-freedom. P-values from this method are adjusted by the Tukey HSD test for a family of 4 estimates (genotype). Data are back-transformed EMM +/− 95% parametric confidence intervals computed from the fixed effects estimated by the linear mixed model.

*Dynamic phase synchronization values* or phase sync slopes for each site-pair were computed differently for individuals and for genotype groups. For heatmaps and PCA, within-individual slopes were computed using a LMM (REML) with numeric activity value, random effect of session, and extracted using lstrends. The model formula was: `asin(sqrt(MRL) ∼ activityNum_lmer * idNum * freqband * compSites + (1|session)`. For statistical analysis, group-wise slopes relating transformed MRL as a function of activity were estimated similarly to before but using LMM (REML) with a numeric activity value, random slopes of session and individuals, and extracted using lstrends. The model formula was: `asin(sqrt(MRL)) ∼ activityNum_lmer * geno * freqband * compSites + (1|session) + (1|idNum)`. Statistical inference on group-wise differences in slope were performed using Tukey contrasts as described previously. Data are back-transformed EMM +/− 95% parametric confidence intervals computed from the fixed effects estimated by the LMM as described.

## ACKNOWLEDGEMENTS

We thank Dr. Huda Zoghbi for support of the early stages of the work and review of the manuscript, and Drs. Jeffrey Neul, Hongwei Dong and Syd Cash for review of the manuscript. We thank the Intellectual and Development Disabilities Research Center at Baylor College of Medicine for use of the Preclinical and Clinical Outcomes and Circuit Analysis and Modulation Core Facilities (formerly Neurobehavioral and Neurophysiology Cores) supported by NIH NICHD *Eunice Kennedy Shriver* grant number P50HD103555 (JT, RCS), and Texas Children’s Hospital. This work was supported by the U.S. NIH NICHD *Eunice Kennedy Shriver* grant numbers R01HD083181 (RCS), NIH Office of the Director grant number DP5OD009134 (RCS), NIH NIGMS grant number T32GM007526 (SS), and NIH NINDS grant numbers T32NS043124 (SS), K08NS118107 (CMM) and R25NS070680 (CMM); the CURE Taking Flight Award (CMM); and the Stedman West Endowed Fund in Neurological Research (RCS). The content is solely the responsibility of the authors and does not necessarily represent the official views of the National Institutes of Health.

## FOOTNOTES

## Author Contributions

Conceptualization: CMM, RCS; Methodology: CMM, RCS; Software: CMM; Validation: CMM, RCS; Formal analysis: CMM, SS, SH, DRC, JT, RCS; Investigation: CMM, SS, SH, DRC, AC, ZW, AJL, YS, JT, RCS; Resources: CMM, JT, RCS; Data curation: CMM, SS, SH, DRC, JT, RCS; Writing - original draft: CMM; Writing - review & editing: CMM, SS, SH, DRC, AC, ZW, AJL, YS, JT, RCS; Visualization: CMM, RCS; Supervision: CMM, RCS; Project administration: CMM, RCS; Funding acquisition: CMM, RCS.

## Data and materials availability

The data that support the findings of the study are available from the corresponding authors upon reasonable request.

