## Supplementary Information for "Counter-balancing X-linked *Mecp2* hypofunction by hyperfunction ameliorates disease features in a model of Rett syndrome: implications for genetic therapies"

#### AFFILIATIONS

[†] Houston Methodist Research Institute, Department of Neurosurgery and Center for Neuroregeneration, Houston, TX, USA.

[‡] College of Life and Health Science, Northeastern University, Shenyang, China.
[§] Medical Scientist Training Program, University of Pennsylvania Perelman School of Medicine, Philadelphia, PA, USA.
[‖] Association of University Centers on Disabilities, Silver Spring, MD, USA.

Figure 1. Additional neurobehavioral phenotyping of Mecp2 counter-balanced mice.
Figure 2. Method of motion estimation.
Figure 3. Contributions of PCA from within-region and between-region analyses of LFP
Figure 4. Quantitative PCR genotyping assay for differentiating animals harboring single human MECP2 transgenes from double transgenic animals.
Figure 5. Positions of depth electrodes used in electrophysiology studies.
Figure 6. Relative power stratified by site, frequency band, and activity state in Mecp2 mutant animals.
Figure 7. Phase synchronization stratified by site-pair, frequency band, and activity state in Mecp2 mutant animals.

### Supplementary Methods

#### Quantitative PCR (qPCR) copy number genotyping assay to distinguish TG alleles.

Tail biopsy samples were obtained from test progeny and genomic DNA was extracted according to the manufacturer's protocol (Gentra® Puregene® Mouse Tail Kit, Qiagen, USA). Endpoint PCR was used as described to confirm the N allele[1]; as expected from the mating scheme design to obtain test progeny, all female mice carried the paternally inherited N allele. Endpoint PCR was used as described to detect mice that were positive for harboring a TG allele[2,3]. Given that the focus of the current study was a comparison of N mice with inheritance of either the TG1 or the TG3 allele, and not N mice that inherited both the TG1 and TG3 alleles, we deployed two additional genotyping steps to exclude double transgenic animals.

First, endpoint PCR was used to amplify a specific PCR product from the TG3 but not TG1 allele: forward, 5'-TTGTGTCTTTTGTTGTAAAAATCAAGTGAT-3'; reverse 5'-ACGATCCTCTCCCTATAGTGAGTCGT-3', expected size PCR product, 374 bp. Second, to confirm whether mice harbored a single TG allele (TG1 or TG3) or both TG alleles (TG1 and TG3), we designed a qPCR genotyping assay to differentiate the lines based on relative transgene copy number. To optimize the qPCR method for transgene copy number, a 50 ng/µL master stock containing gDNA from known positive control TG1 and TG3 positive samples was used to create a standard curve comprised of serial 5-fold dilutions. Seven standards, ranging from 10 ng/µL to $6.4 \times 10^{-4}$ ng/µL, were used in qPCR amplification of human *MECP2* and mouse *Pcdhb2.* Human-specific primers targeting genomic 3'UTR of *MECP2* with no mouse cross-reactivity were chosen using Primer-BLAST: forward, 5'-AGGTGAATACGATACAGGGCTT-3'; reverse, 5'GTTGGGAGAGGTGCACTTGG-3'. A region of the mouse gene *Pcdhb2* was selected as a control region: forward, 5'-GGAAATGGAGCAAATCCTGA-3'; reverse, 5'-GTGCAGCGAGTTCCTCTACC-3'. The ratio of human *MECP2* (M) to mouse *Pcdhb2* (P) derived from qPCR signal from gDNA samples corresponded with the presence of TG1, TG3 or TG1;TG3 alleles. Based on optimization experiments, the average value of M-to-P expressed as a ratio denoted either the presence of the TG1 allele alone ($M/P < 0.4$), TG3 allele alone ($M/P = 0.5 - 0.85$), or both the TG1 and TG3 alleles ($M/P > 0.87$). An example of qPCR copy number results is shown in **Supplementary Fig. 4**.

Quantitative PCR analysis of each gDNA sample for confirmatory genotype assignment was then conducted in triplicate and compared with the optimized copy number threshold to detect TG1 alone, TG3 alone or TG1 and TG3. Care was taken in handling gDNA to minimize variability, with limited freeze-thaw of samples across testing. Reactions were made using 2X SYBR Green Supermix (Bio-Rad, USA) with either 10 µM of primer set for detection of *hMECP2* or 20 µM *mPcdhb2* diluted in DNA hydration buffer (Qiagen, USA) containing 30 ng/µL salmon sperm DNA (Thermo Fisher Scientific, USA). At least two independent TG1 and TG3 biological replicates and a no template control were analyzed in triplicate alongside each set of target samples as additional positive and negative controls, respectively.

**Western blot.**

Tissue for Western blot analysis was obtained from whole mouse brain. In brief, brain was dissected on ice and immediately frozen in liquid nitrogen and stored at -80degC. Frozen tissue was homogenized directly in lysis buffer (0.1 M Tris HCl pH 7.5, 2%SDS, with protease inhibitor (Sigma P8340)) using a 2ml dounce homogenizer in ice slurry, 20 stokes with loose pestle A, and 20 stokes with tight pestle B, followed by sonication, and 10 passages through 27G needle. Finally, samples were centrifuged x 10 minute at 15,000 RPM in a refrigerated microcentrifuge at 4 °C. Supernatants were transferred to clean Eppendorf tubes and frozen at -20degC. Primary antibodies included rabbit anti-MECP2 (Sigma, #M7443, 1:1000), and rabbit anti-histone 3 (H3) antibody (Millipore #07-690, 1:45000). Signal was detected using HRP-conjugated secondary antibodies (Amersham ECL Donkey Anti-rabbit IgG HRP (#NA934), 1:5000; Amersham ECL Sheep Anti-mouse IgG HRP (#NA931), 1:5000) and developed using Pierce ECL reagents on an Image Quant LAS 4000. Quantification was performed using ImageJ.

**Immunofluorescence and quantification.**

As previously described,[4,5] brains were fixed by transcardial perfusion of phosphate buffered saline (PBS) followed by 4% paraformaldehyde (PFA) in PBS. The brains were then postfixed in PFA for 3 hours, cryoprotected in 30% sucrose in PBS, embedded in optimal cutting temperature medium (O.C.T), and subsequently frozen. Free-floating coronal sections at 50 µm thickness were obtained using a Leica CM3050S cryostat. Matched sections were selected and washed in PBS for antibody incubation as floating sections. Sections were washed in PBS and subsequently permeabilized in 0.4% Triton-X 100 in PBS (PBST). The sections were then blocked for two hours in blocking buffer containing 5% normal donkey serum, 1% bovine serum albumin, and 0.1% glycine in PBST. After blocking, the sections were then incubated in primary antibody (rabbit anti-MeCP2, Cell Signaling; mouse anti-NeuN, Cell Signaling) x 24hrs at 4°C. The sections were subsequently washed in PBST, then incubated in Alexa Fluor 488-conjugated Donkey Anti-Mouse IgG (1:500; Jackson ImmunoResearch) and Alexa Fluor 555 Anti-Rabbit IgG secondary antibody (1:500) in blocking buffer for 4 hours at room temperature. The sections were then washed in PBST, incubated in DAPI in PBST (1:10,000) for 15 minutes, and washed in PBST again. The sections were mounted onto slides using Prolong Gold Antifade Mounting Medium (ThermoFisher Scientific, USA). Slides were imaged using a Zeiss LSM 880 confocal microscope with a 63x objective. Z-stack images were captured with identical image acquisition settings, and consistent regions of interest were imaged using anatomical landmarks based on The Mouse Brain in Stereotaxic Coordinates, 3rd edition (Franklin and Paxinos, 2007). Three sections per animal were imaged, and maximum-intensity projections were prepared using the Zeiss Zen 2 software.
*Quantification of MECP2 immunofluorescence.* Automated quantification of MeCP2 fluorescence signal intensity in individual neurons was accomplished using Bitplane Imaris 64x version 7.7.1. Using the Imaris surface tool, surfaces were created to encapsulate individual nuclei based on a threshold mean DAPI

fluorescence intensity and a minimum 3D volume. Within the NeuN-positive surfaces (which had a threshold mean NeuN fluorescence intensity), the mean MeCP2 signal intensity was calculated. The MeCP2 fluorescence intensities per each NeuN-positive cell were then plotted together. Four images were analyzed per genotype, and n=3 animals per genotype were used. Over 2000 individual cells were quantified, using consistent intensity thresholds.

**Transcriptional analysis.**

Hypothalamus was dissected from adult mice as previously described[6]. RNA was isolated from Trizol, processed using RNeasy Cleanup Kit (Qiagen) and cDNA synthesis performed using SuperScript III RT (Invitrogen). Quantitative real-time PCR was performed using custom primers and Quanta PerfeCTa SYBR Green Fast Mix on a CFX96 Real-Time PCR detection system with C1000 thermal cycler (Biorad). Results were quantified using the standard curve method and normalized to detected levels of S16. Statistical significance was calculated using one-way ANOVA (genotype) and Fisher's LSD post-hoc test.

**Supplementary Figures**

**Figure 1.** Additional neurobehavioral phenotyping of Mecp2 counter-balanced mice.

**Figure 2.** Method of motion estimation.

**Figure 3.** Contributions of PCA from within-region and between-region analyses of LFP.

**Figure 4.** Quantitative PCR genotyping assay for differentiating animals harboring single human MECP2 transgenes from double transgenic animals.

**Figure 5.** Positions of depth electrodes used in electrophysiology studies.

**Figure 6.** Relative power stratified by site, frequency band, and activity state in Mecp2 mutant animals.

**Figure 7.** Phase synchronization stratified by site-pair, frequency band, and activity state in Mecp2 mutant animals.

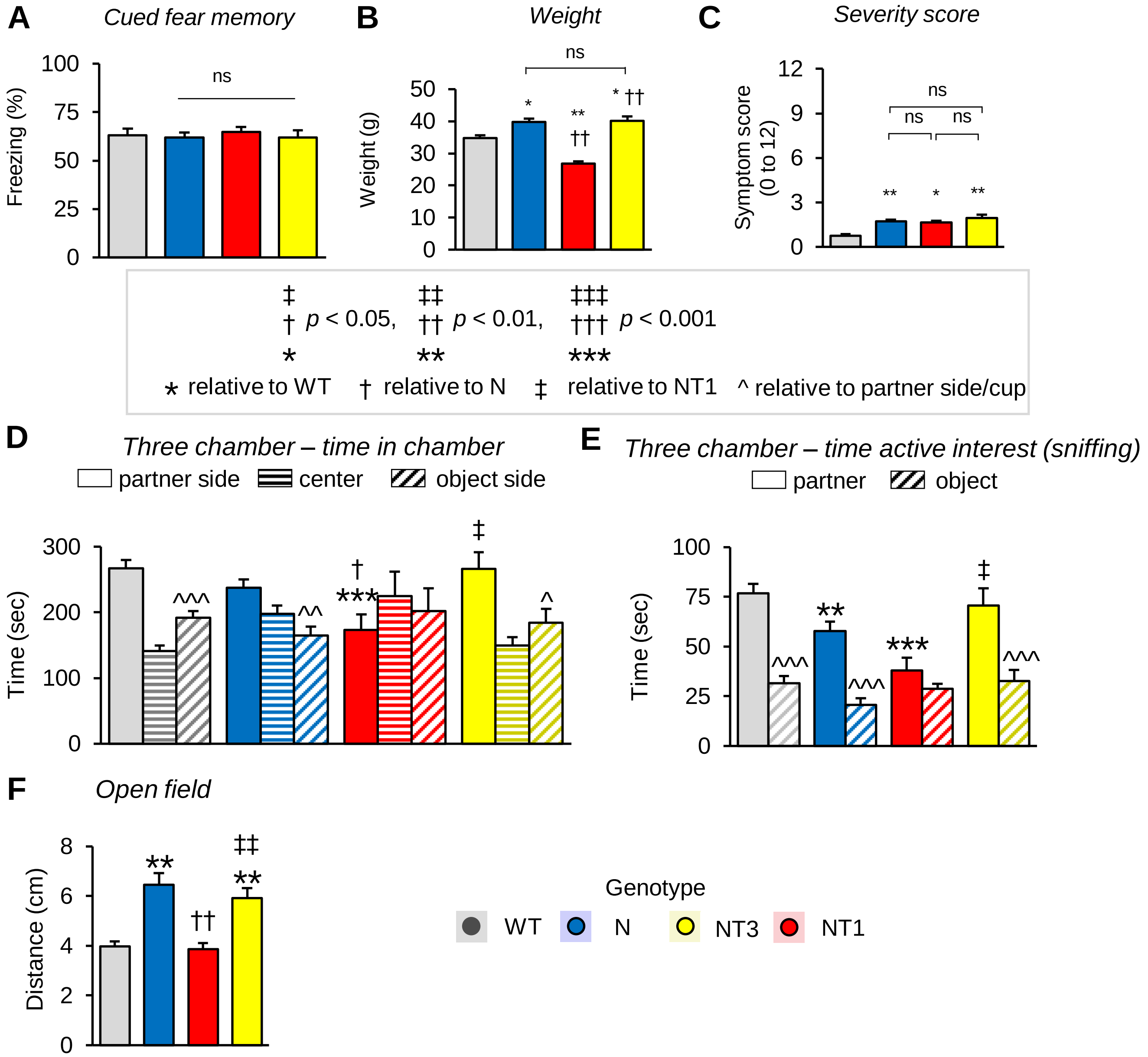

**Supplementary Figure 1. Additional neurobehavioral phenotyping of Mecp2 counter-balanced mice.** (**A**) Cued fear. (**B**) Weight. (**C**) Symptom score. (**D-E**) Three-chamber sociability test showing time in chamber (D) and time spent actively sniffing (E). (**F**) Open field activity. For A-F, data represent the mean ± sem, N=24-31 mice per genotype. Symbols (*, †, ‡) represent differences relative to WT, N, or NT1 respectively. For each symbol, a single symbol denotes $p < 0.05$; double symbol denotes $p < 0.01$; and triple symbol denotes $p < 0.001$.

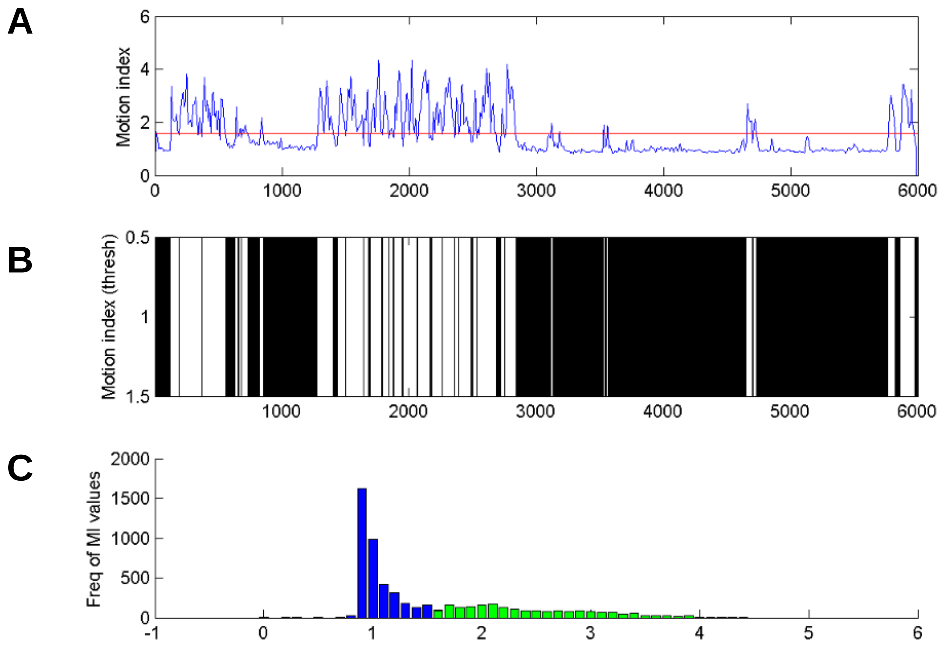

**Supplementary Figure 2. Method of motion estimation.**
**(A-B)** Representative time-series from individual mouse showing motion indices (MIs) representing frame-wise intensity-differences following preprocessing (**A**) and binarized data based on threshold >= 1.6 (**B**). X-axis is time (seconds). (**C**) Histogram of MI values demonstrates MI values < 1.6 (*blue*) correspond with inactivity, while MI >= 1.6 correspond with activity (*green).*

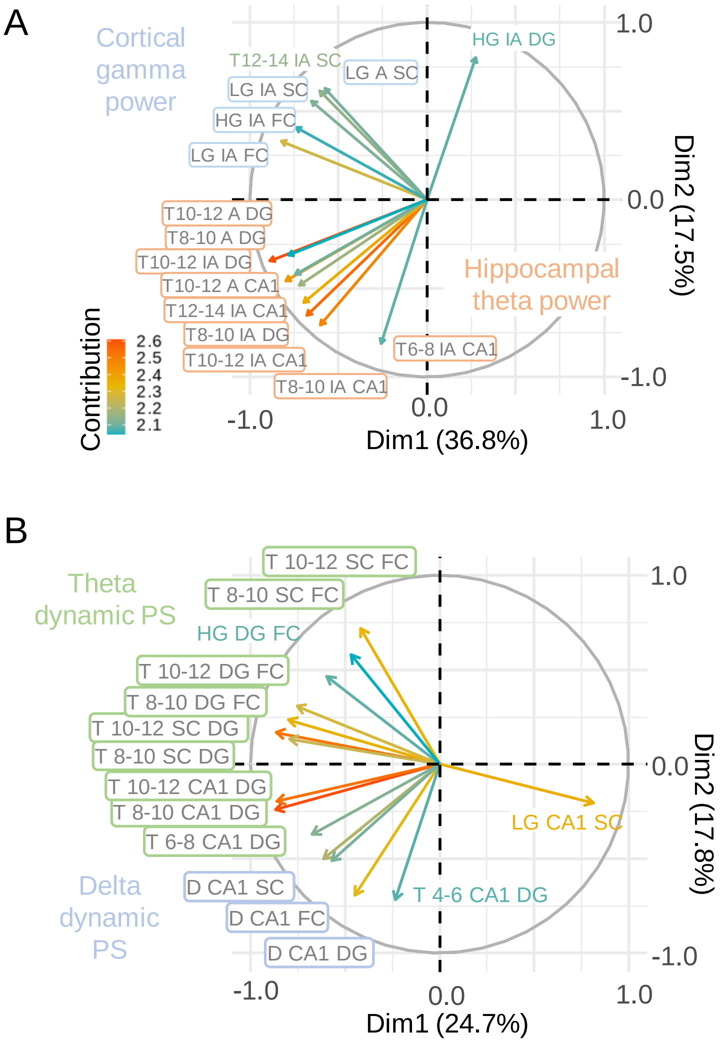

**Supplementary Figure 3. Contributions of PCA from within-region and between-region analyses of LFP.** (A) Top 15 contributing factors from PCA of relative band-passed power from Figure 3. (B) Top 15 contributing factors from PCA of dynamic phase synchronization values from Figure 4.

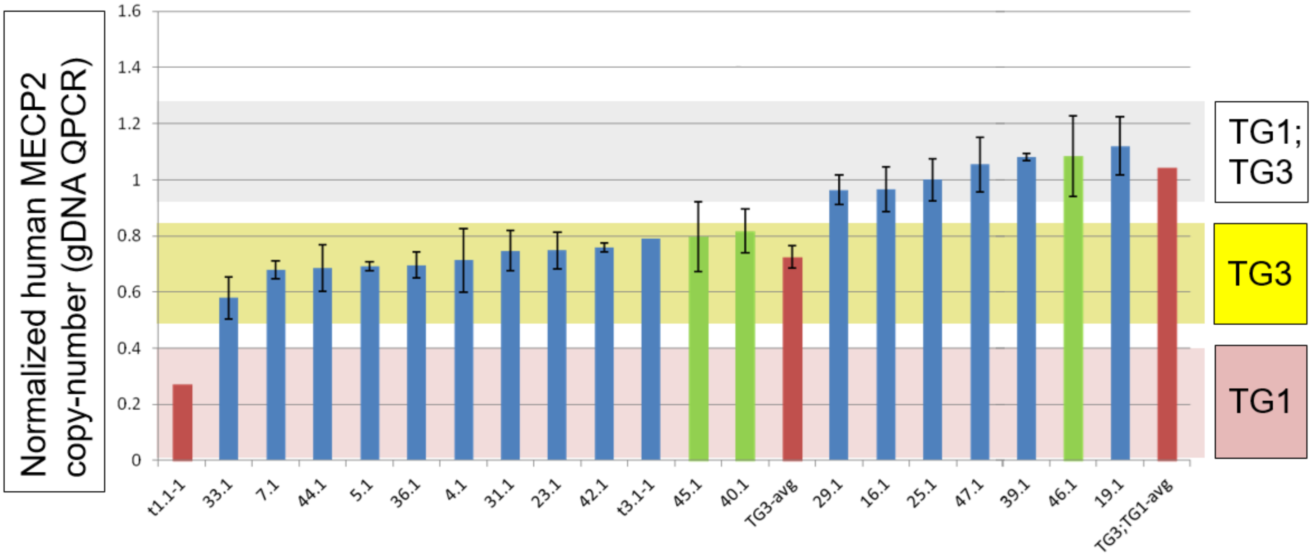

**Supplementary Figure 4. Quantitative PCR genotyping assay for differentiating animals harboring single human MECP2 transgenes from double transgenic animals.** Data is the ratio of human *MECP2* copy number to autosomal mouse reference gene, *pcdh2b.* Bars indicate mean +/- s.e.m from N=3 technical replicates of individual animals. Animals harboring the TG3 allele alone were observed to have M/P = 0.5 - 0.85, while animals with both TG1 and TG3 alleles have M/P > 0.87. Double transgenic animals were not used in our experiments.See Methods for additional details. Blue bars show individual test animals, green bars show positive controls, red bars indicate mean +/- s.e.m. values for indicated sample type.

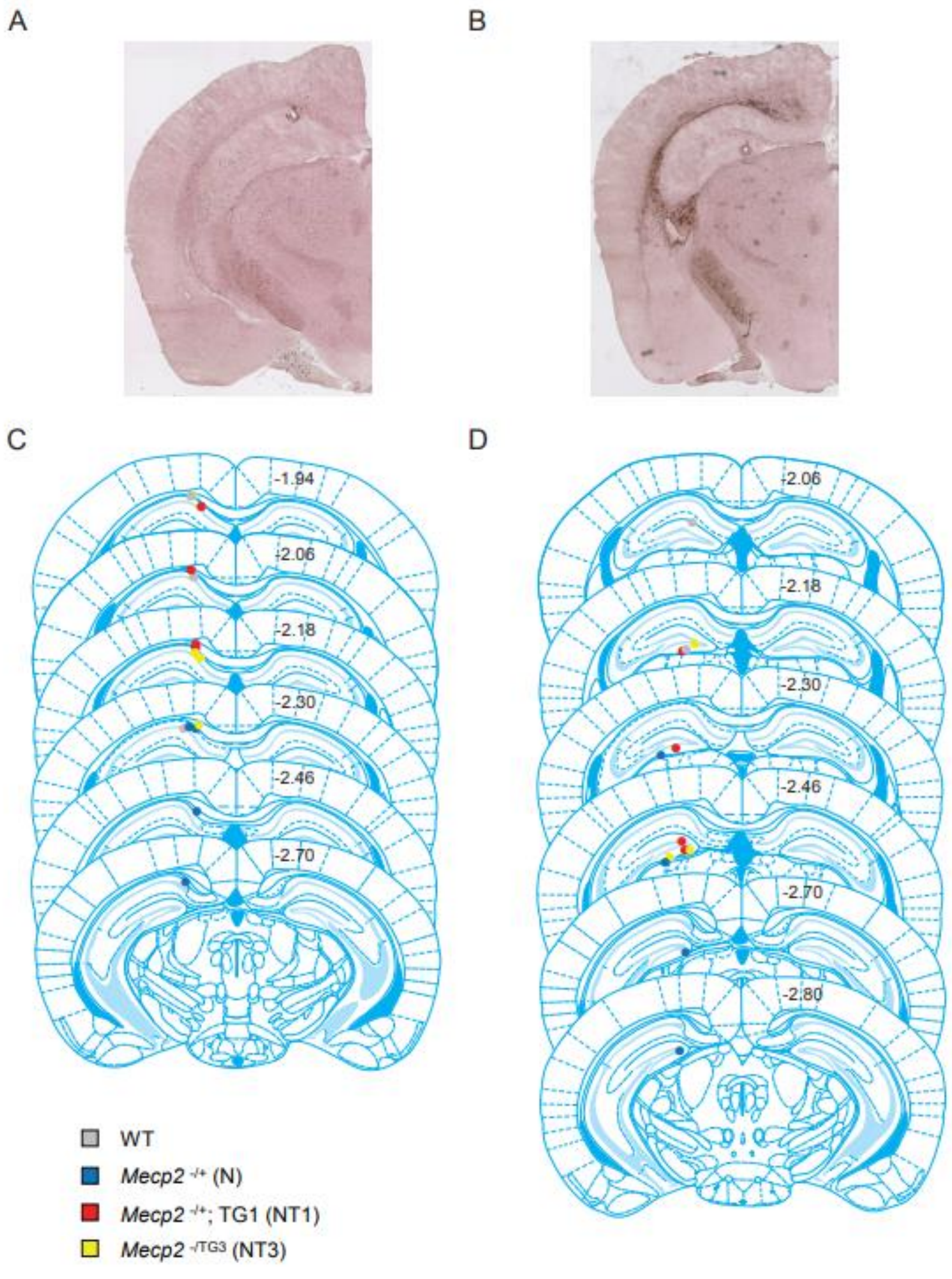

**Supplementary Figure 5. Positions of depth electrodes used in electrophysiology studies. (A-B**) Representative coronal sections demonstrating position of electrolytic lesion associated with depth electrode in hippocampal CA1 (A) and dentate gyrus (B). (**C-D**) Schematic of electrode lesions from all animals (N=16) in hippocampal CA1 (C) and dentate gyrus (D). Atlas drawings adapted from Paxinos et al.

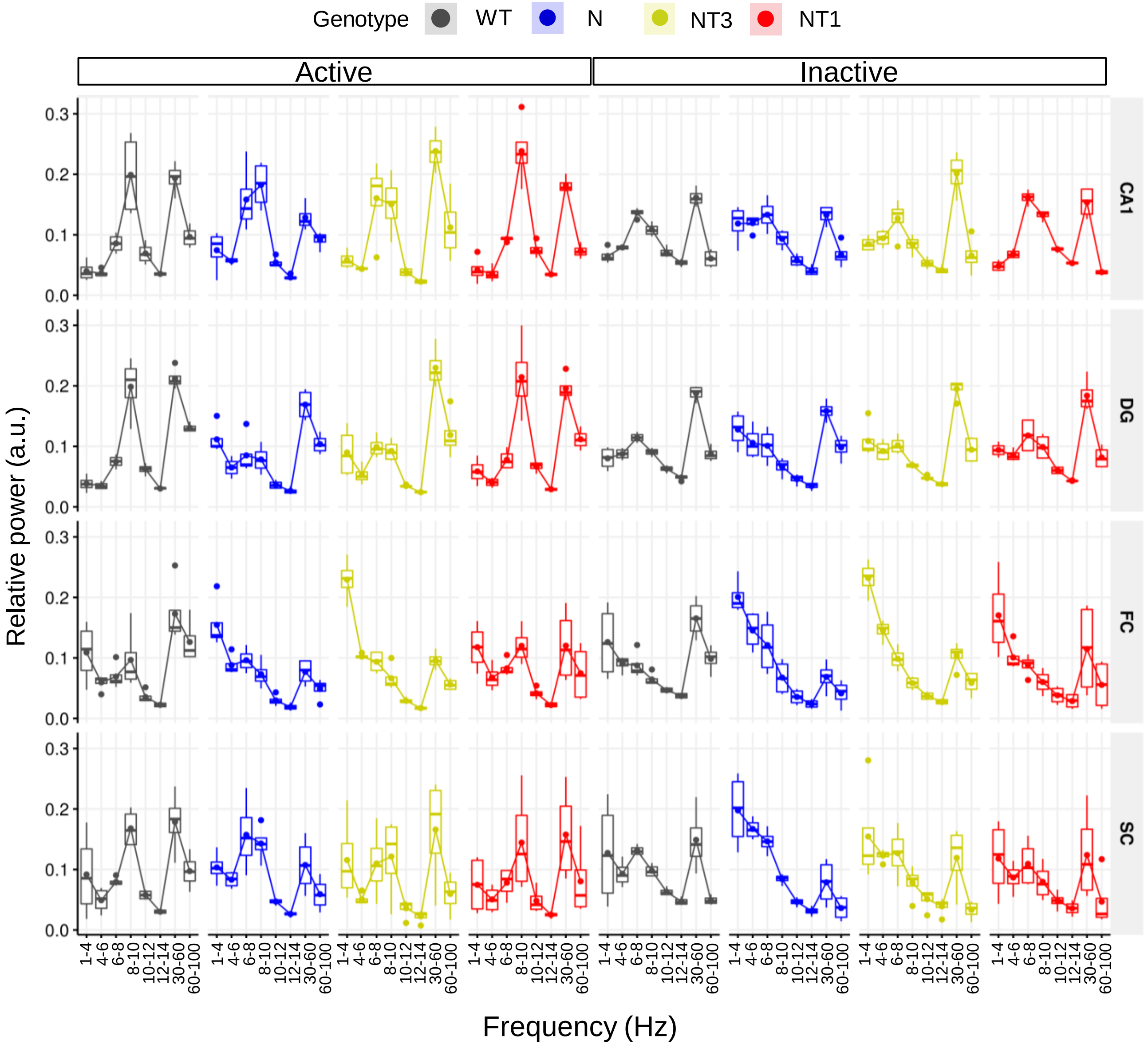

**Supplementary Figure 6. Relative power stratified by site, frequency band, and activity state in Mecp2 mutant animals.** Data are box-and-whisker plots of measurements from each of N=4 animals per genotype, averaged over N=4 sessions (1-2 hours per session). X-axis represents eight frequency bands, y-axis is relative band-specific power during activity (activeAll)) or inactivity (inactiveAll) within each brain region (*rows*, CA1, DG, FC, and SC).

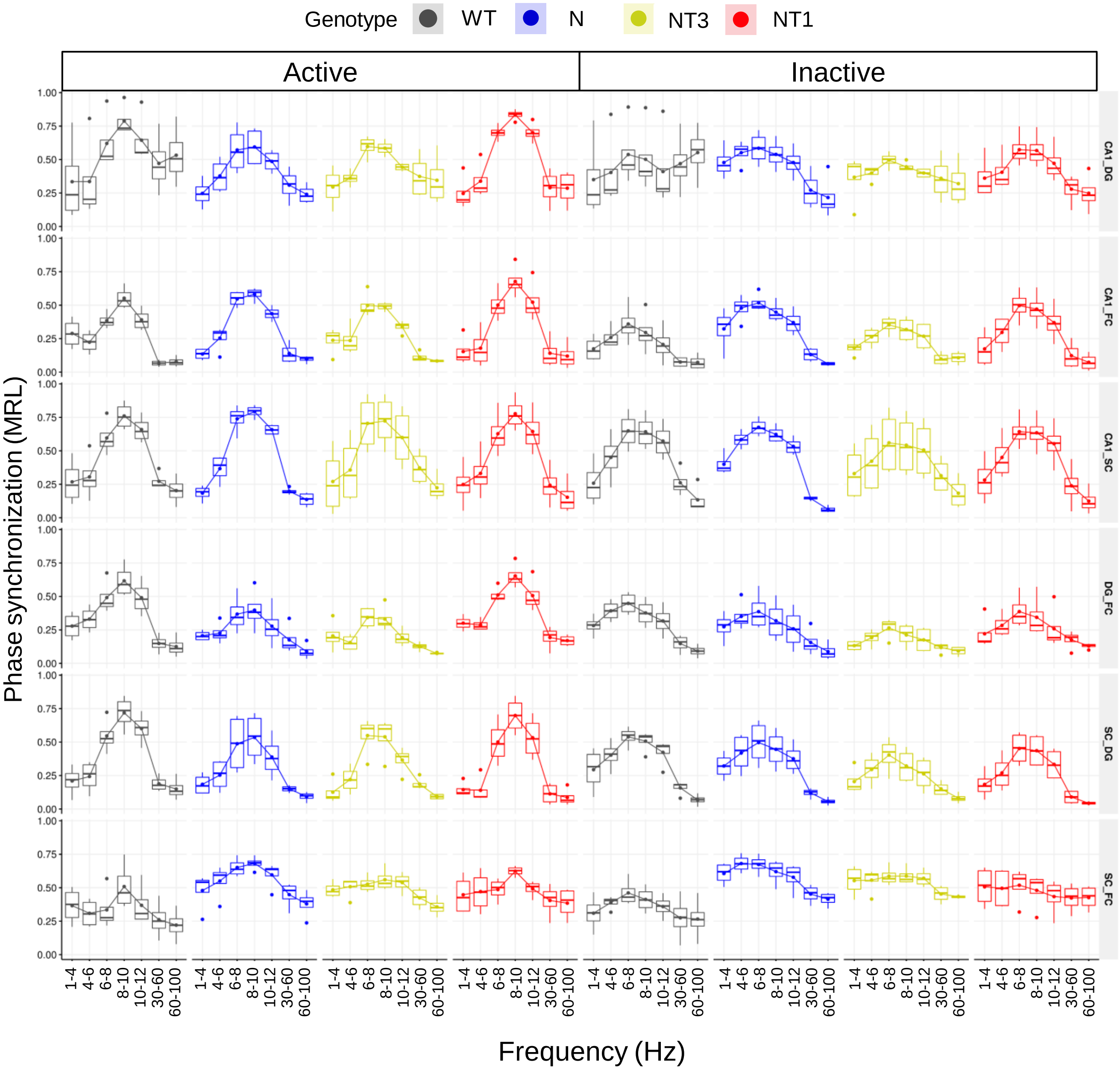

**Supplementary Figure 7. Phase synchronization stratified by site-pair, frequency band, and activity state in Mecp2 mutant animals.** Data are box-and-whisker plots of measurements from each of N=4 animals per genotype, averaged over N=4 sessions (1-2 hours per session). X-axis represents sevene frequency bands, y-axis is band-specific phase synchronization during activity or inactivity within each brain region pair (*rows).* Sub-bands within the theta range are labelled according to their corresponding frequency range. The theta band from 12-14Hz did not show major differences in preliminary analyses and was therefore excluded for simplicity.
